# Shortened TDP43 isoforms upregulated by neuronal hyperactivity drive TDP43 pathology in ALS

**DOI:** 10.1101/648477

**Authors:** Kaitlin Weskamp, Elizabeth M. Tank, Roberto Miguez, Jonathon P. McBride, Nicolás B. Gómez, Matthew White, Ziqiang Lin, Carmen Moreno Gonzalez, Andrea Serio, Jemeen Sreedharan, Sami J. Barmada

**Author notes:** To whom correspondence should be addressed: Sami Barmada, University of Michigan, Department of Neurology, 109 Zina Pitcher Place, BSRB 5015, Ann Arbor, MI 48109.

## Abstract

Cortical hyperexcitability and mislocalization of the RNA-binding protein TDP43 are highly-conserved features in amyotrophic lateral sclerosis (ALS). Nevertheless, the relationship between these phenomena remains poorly defined. Here, we showed that hyperexcitability recapitulates TDP43 pathology by upregulating shortened (s) TDP43 splice isoforms. These truncated isoforms accumulated in the cytoplasm and formed insoluble inclusions that sequestered full-length TDP43 via preserved N-terminal interactions. Consistent with these findings, sTDP43 overexpression was toxic to mammalian neurons, suggesting neurodegeneration arising from complementary gain- and loss-of-function mechanisms. In humans and mice, sTDP43 transcripts were enriched in vulnerable motor neurons, and we observed a striking accumulation of sTDP43 within neurons and glia of ALS patients. Collectively, these studies uncover a pathogenic role for alternative TDP43 isoforms in ALS, and implicate sTDP43 as a key contributor to the susceptibility of motor neurons in this disorder.

## Introduction

Amyotrophic lateral sclerosis (ALS) is a neurodegenerative disorder in which the progressive loss of motor neurons results in paralysis and respiratory failure (1). There is no disease-modifying therapy for ALS, and its heterogeneous biochemical, genetic, and clinical features complicate the identification of therapeutic targets. However, the cytoplasmic mislocalization and accumulation of TDP43 (TAR DNA-binding protein of 43 kD), a nuclear RNA-binding protein integrally involved in RNA metabolism, is observed in >90% of individuals with ALS (2). Moreover, while mutations in the gene encoding TDP43 (*TARDBP*) only account for 2-5% of ALS cases, mutations in several other ALS-associated genes including *C9ORF72* (3), *ANG* (4), *TBK1* (5), *PFN1* (6), *UBQLN2* (7), *VCP* (8), and *HNRNPA2B1* (9) result in TDP43 pathology.

TDP43 is an essential protein involved in several RNA processing events, including splicing, translation, and degradation. In keeping with these fundamental functions, TDP43 levels and localization are tightly regulated and critical for cell health. TDP43 knockout animals exhibit neurodegeneration and behavioral deficits (10–13), while TDP43 overexpression results in neurodegeneration in primary neuron (14, 15), mouse (16, 17), rat (18, 19), Drosophila (20, 21), zebrafish (22, 23), and primate models (24, 25). Furthermore, mislocalization of TDP43 to the cytoplasm is sufficient to drive cell death (14). Taken together, this suggests that even small changes to TDP43 levels and localization are highly predictive of neurodegeneration.

Hyperexcitability, or an increase in neuronal activity, is also a conserved feature in both familial and sporadic ALS (26). Cortical hyperexcitability precedes symptom onset in some cases (26), and the degree of motor neuron excitability is a strong predictor of disease progression (27, 28). Such hyperexcitability arises from a loss of cortical inhibition (26, 29–33) in combination with intrinsic differences in channel expression, content, and activity within motor neurons themselves (26, 28, 34, 35). Emphasizing the contribution of hyperexcitability to disease, riluzole—one of two available therapies for ALS—is a sodium channel antagonist that partially rescues hyperexcitability (36). Animal models of ALS recapitulate key features of hyperexcitability (37–39), including an increase in motor neuron activity that precedes the onset of motor deficits (37, 39, 40) and reduced activity following treatment with riluzole (41). Hyperexcitability is also observed in iPSC-based ALS models (42, 43), though other reports suggest that it may be a transient or developmental phenomenon (43, 44).

Despite the prevalence of both TDP43 pathology and hyperexcitability in ALS, the relationship between these phenomena remains poorly defined. Here, we utilize an iPSC-derived neuron (iNeuron) model system to demonstrate that hyperexcitability drives TDP43 pathology characteristic of ALS via the upregulation of atypical, shortened TDP43 isoforms. Using multiple model systems and human post-mortem material, we show that these unusual isoforms are exported from the nucleus, form insoluble cytoplasmic inclusions, are neurotoxic, and are enriched in ALS patient tissue, thereby directly implicating alternative TDP43 isoforms in ALS pathogenesis.

## Results

### TDP43 is regulated by neuronal activity

To investigate disease mechanisms related to hyperexcitability in human neurons, we established an induced pluripotent stem cell (iPSC) derived neuron (iNeuron) model. Transcription activator-like endonucleases (TALENs) specific for the *CLYBL* safe harbor locus were used to introduce the transcription factors Neurogenin 1 and 2 (Ngn1-2) under a doxycycline (dox)-inducible promoter (Figure 1A). Expression of Ngn1-2 is sufficient to drive the rapid differentiation of iPSCs into iNeurons that display immunocytochemical and electrophysiological properties of glutamatergic, excitatory forebrain-like neurons (45–47) (Figure 1B). Consistent with this, within 2 weeks of dox addition iNeurons adopt a neuronal morphology and stain positive for the neuronal markers VGLUT1 and TUJ1 (Figure 1C). We further validated the maturity of neurons differentiated in this manner using an iPSC line that stably expresses the fluorescent calcium indicator gCaMP6f in addition to dox-inducible Ngn1-2 (48). Because time-dependent changes in gCaMP6f fluorescence correlate with action potentials, we monitored neuronal activity indirectly and non-invasively in iNeurons by fluorescence microscopy. Two to three weeks following dox addition, iNeurons displayed a low level of spontaneous activity that was significantly increased with bath application of the neurotransmitter glutamate or the potassium channel blocker tetraethylammonium (TEA; Figure 1D-F). Conversely, activity was inhibited by application of the sodium channel blocker tetrodotoxin (TTX). Though glutamate dramatically increased neuronal activity, it proved to be toxic even at low doses (data not shown). In comparison, iNeurons treated with TEA showed a smaller, sustained increase in activity without significant cell death (Figure 1G). Thus, TEA and TTX were utilized in future studies of activity-dependent TDP43 regulation.

**Figure 1.**
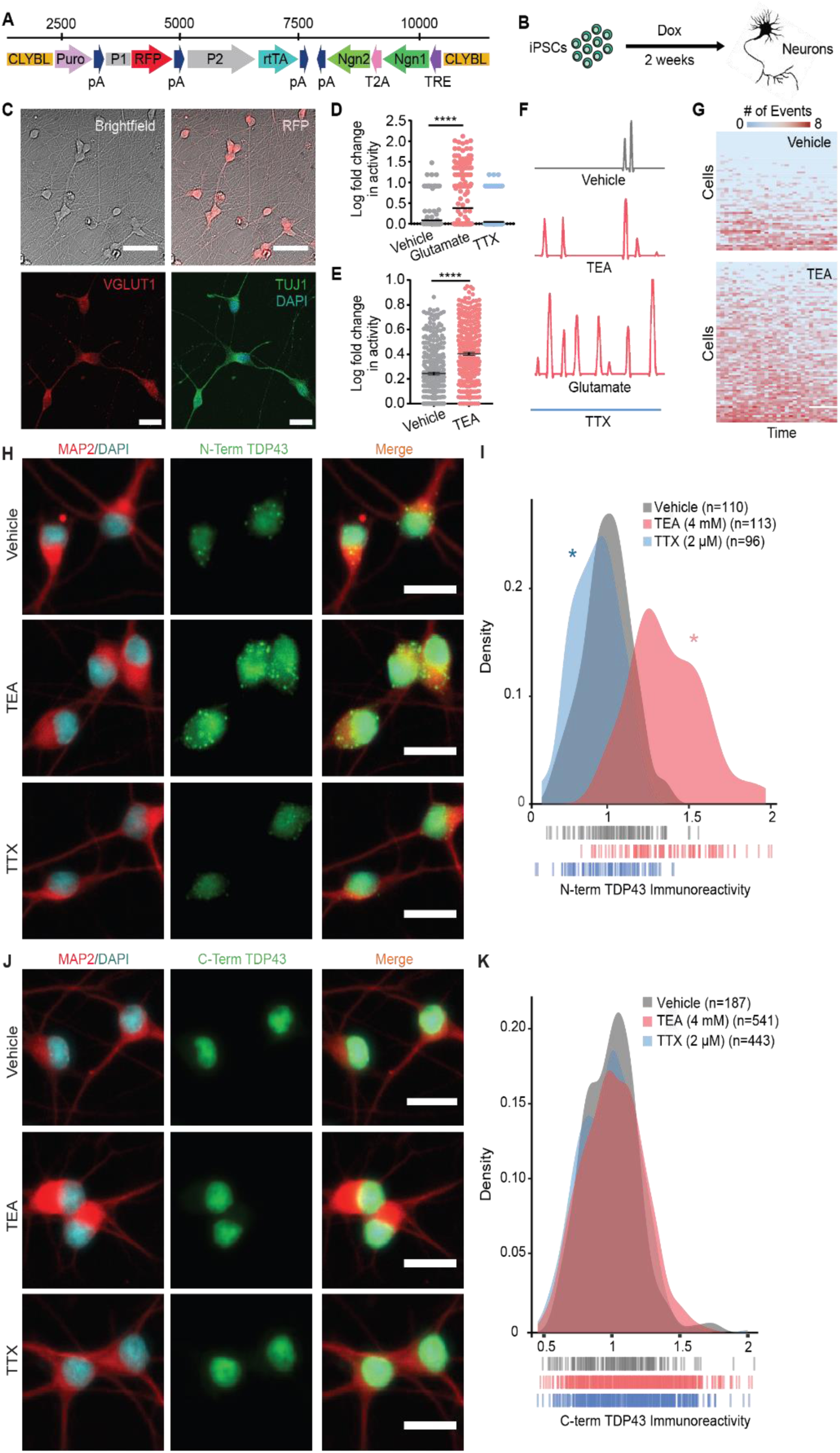
Hyperexcitability drives TDP43 accumulation in human iNeurons. (**A**) Schematic of the cassette used to integrate Ngn1 and Ngn2 into the *CLYBL* safe harbor locus under a doxycycline-inducible (Tet-on) promoter. CLYBL, targeting sequence; Puro, puromycin resistance gene; pA, poly-A tail; P1, P2, promoters; RFP, mCherry; rtTA, reverse tetracycline-controlled transactivator; Ngn1, 2, Neurogenin1 and 2; T2A, self-cleaving peptide; TRE, tetracycline response element. (**B**) Timeline depicting the differentiation of iPSCs into forebrain-like neurons within 2w of doxycycline addition. (**C**) The resultant neurons are RFP-positive and express the neuronal markers VGLUT1 and TUJ1. (**D**) Spontaneous neuronal activity visualized by the Ca^2+^ reporter gCaMP6f at 2w. Activity was pharmacologically modulated with bath application of glutamate or TTX. Vehicle n=257, Glutamate n=327, TTX n=403, stratified among 3 replicates; ****p<0.0001, one-way ANOVA with Dunnett’s post-test. (**E**) Treatment with TEA significantly increased neuronal activity. Vehicle n=312, TEA n=369, stratified among 3 replicates, ****p<0.0001, two-tailed t-test. (**F**) Example traces depicting changes in gCaMP6f fluorescence for each condition. (**G**) Heat maps depicting global changes in activity. Each row represents one neuron, and each column represents a 20s observation window. Thirty intervals were collected over a 12h period. Box color indicates the relative firing rate of each cell at each timepoint ranging from low (blue) to high (red). (**H**) N-terminal TDP43 immunoreactivity was increased in TEA-treated iNeurons and decreased in TTX-treated iNeurons. (**I**) Density plot depicting the change in TDP43 immunoreactivity between conditions. Vehicle n=110, TEA n=113, TTX n=96, 2 replicates, dashes indicate single neurons, *p<0.05, Kolmogorov-Smirnov test. (**J**) No change in TDP43 abundance was detected using an antibody directed against the C-terminus. (**K**) Density plot depicting the change in C-terminal TDP43 immunoreactivity between conditions. Vehicle n=187, TEA n=541, TTX n=443, 2 replicates, dashes indicate single neurons, not significant by the Kolmogorov-Smirnov test. Scale bars in (**C**), 50 µm top, 20 µm bottom. Scale bars in (**H**), (**J**), 20 µm.

To explore a potential connection between hyperexcitability and TDP43 pathology, we pharmacologically stimulated or blocked activity in iNeuron cultures and then examined changes in TDP43 levels via immunocytochemistry (ICC) using an antibody directed against the TDP43 N-terminus. To quantitatively gauge differences in neuronal TDP43, we utilized MAP2 staining to generate cellular regions of interest (ROIs), and measured TDP43 immunoreactivity within individual neurons. TEA-treated iNeurons showed a significant increase in TDP43 immunoreactivity while TTX-treated iNeurons exhibited a reduction, suggesting a bidirectional relationship between TDP43 abundance and neuronal activity (Figure 1H, I). An analogous relationship was observed in rodent primary mixed cortical neurons treated with glutamate or the GABA receptor antagonist bicuculline (Supplemental Figure 1A).

Unexpectedly, when we repeated these studies using an antibody directed against the TDP43 C-terminus, we failed to identify significant activity-dependent changes in protein abundance (Figure 1J, K), and also noted prominent differences in subcellular TDP43 distribution identified by each antibody (Figure 1H, J). Immunostaining with N-terminal antibodies revealed punctate, cytoplasmic TDP43 superimposed upon nuclear TDP43 in both iNeurons (Figure 1H) and rodent primary mixed cortical neurons treated with bicuculline (Supplemental Figure 1B). However, only nuclear TDP43 was detectable using C-terminal TDP43 antibodies (Figure 1J). A survey of commercially available antibodies with known epitopes revealed similar trends in localization: antibodies that recognize the TDP43 N-terminus are more likely to display nuclear and cytoplasmic staining patterns (49–51), while antibodies specific to the C-terminus primarily show nuclear TDP43 (52, 53).

Given the variability in antibody specificity and potential difficulties in reproducing results using different antibodies (54, 55), we validated our findings by fluorescently-labeling native TDP43 in iPSCs using CRISPR/Cas9 genome engineering. To minimize off-target effects, we used a dual-nickase approach (56) to fuse the green-fluorescent protein Dendra2 to either the N-terminus (D2-TDP43) or the C-terminus (TDP43-D2) of endogenous TDP43 in human iPSCs (Figure 2A, Supplemental Figure 2). D2-TDP43 and TDP43-D2 iPSCs were differentiated into iNeurons as described before (Figure 2B, C), and neuronal activity was pharmacologically stimulated or blocked using TEA or TTX, respectively. After 48h, we visualized native Dendra2-labeled TDP43 by fluorescence microscopy, noting a bidirectional relationship between D2-TDP43 abundance and neuronal activity (Figure 2D) that was nearly identical to what we observed using antibodies that recognize the TDP43 N-terminus (Figure 1H, I). In comparison, there were no significant activity-dependent changes in TDP43-D2 (Figure 2E), consistent with our inability to detect changes upon staining with antibodies raised against the TDP43 C-terminus (Figure 1J, K). These data provide convincing evidence for TDP43 species harboring the N-but not the C-terminus that are regulated by neuronal activity. Additionally, the distinctive TDP43 distribution patterns revealed by N- and C-terminal reactive antibodies were reflected by the localization of Dendra2-tagged native TDP43: D2-TDP43 appeared both cytoplasmic and nuclear (Figure 2B), while the distribution of TDP43-D2 was limited to the nucleus (Figure 2C).

**Figure 2.**
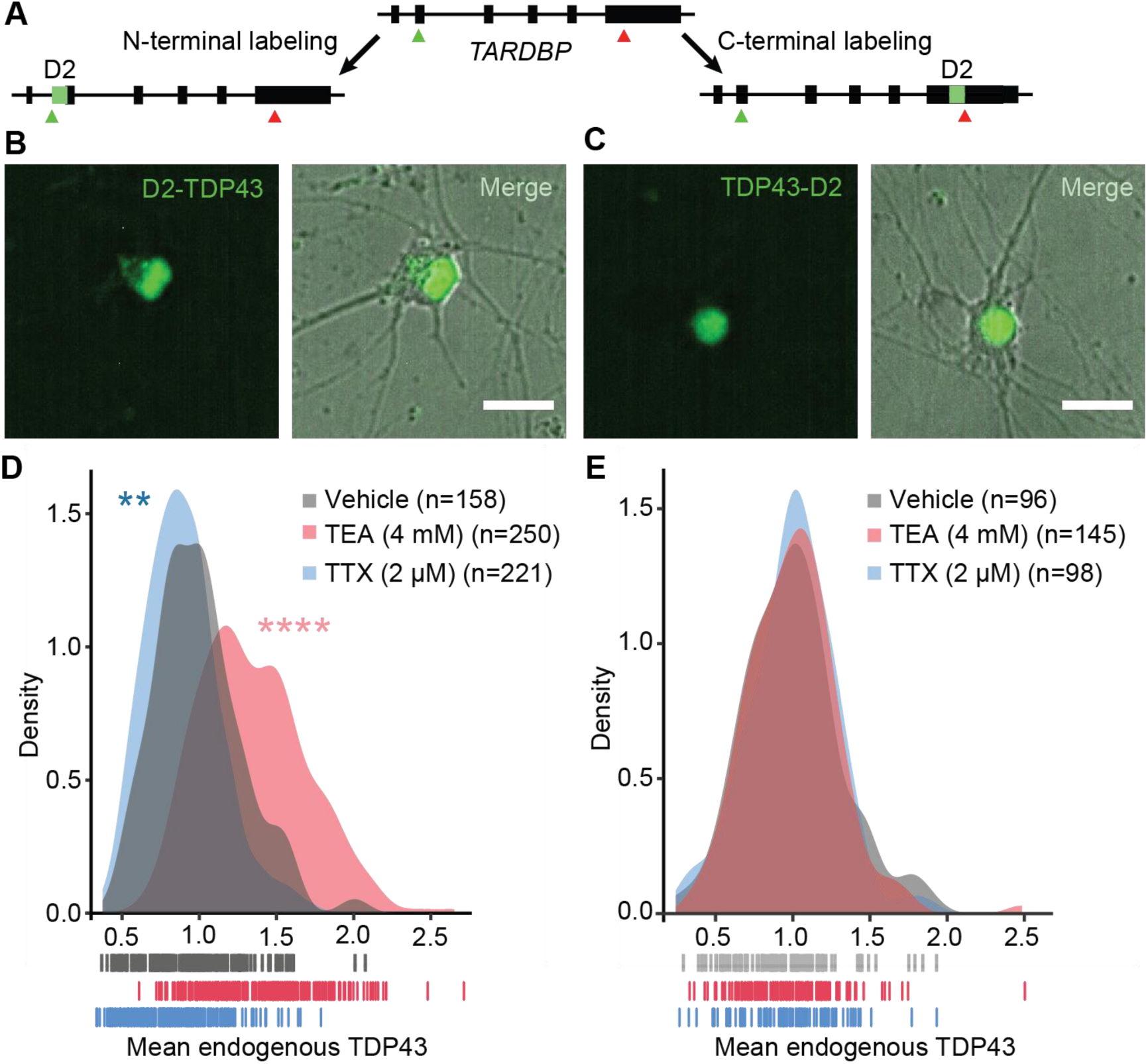
TDP43 species harboring the N-but not the C-terminus are regulated by neuronal activity. (**A**) Strategy for labeling native TDP43 in human iPSC-derived neurons using CRISPR/Cas9. Dendra2 (D2, green) was inserted 3’ to the *TARDBP* start codon (green arrow) or 5’ to the conventional stop codon (red arrow), enabling fluorescent labeling of the TDP43 N- or C-terminus, respectively. In iNeurons, N-terminally tagged TDP43 (**B**, D2-TDP43) appeared both nuclear and cytoplasmic in distribution, while C-terminally tagged TDP43 (**C**, TDP43-D2) was primarily nuclear. (**D**) Density plot depicting the fluorescence intensity of D2-TDP43 upon application of vehicle (n=158), TEA (n=250), or TTX (n=221). (**E**) Density plot depicting the fluorescence intensity of TDP43-D2 with addition of vehicle (n=96), TEA (n=145), or TTX (n=98). In (**D**) and (**E**), dashes indicate individual neurons from 2 replicates, **p<0.01, ****p<0.0001, Kolmogorov-Smirnov test. Scale bars in (**B**) and (**C**), 20 µm.

Collectively, these results suggest that neuronal activity elicits an increase in cytoplasmic TDP43 that lacks a C-terminus. In contrast to what we observed with N-terminal TDP43, there was no reciprocal activity-dependent change in C-terminal TDP43 abundance or localization by immunocytochemistry, and we failed to observe any differences in C-terminally labeled TDP43-D2 upon addition of TEA or TTX, arguing against a cleavage event. However, previous studies demonstrated that neuronal activity regulates the abundance of similar RNA-binding proteins through alternative splicing (57, 58). We therefore considered the possibility that activity gives rise to distinct TDP43 isoforms through alternative splicing.

### Hyperexcitability drives *TARDBP* alternative splicing

Using available RNA-seq data obtained from human cell lines (59), we identified two alternatively spliced *TARDBP* isoforms predicted to encode C-terminally truncated or shortened (s) TDP43 isoforms (Figure 3A). Identical sTDP43 splice isoforms (TDP-S6 and TDP-S7) were detected in previous studies of *TARDBP* splicing (60, 61). Both sTDP43-specific splice donors are located within *TARDBP* exon 6 and differ by only 9 bp; each utilizes an identical splice acceptor within the *TARDBP* 3’ untranslated region (UTR), thereby eliminating the majority of exon 6 (Figure 3B). We designed primers specific for both sTDP43 splice junctions as well as full-length (fl) TDP43 utilizing the canonical termination codon within exon 6, and performed qRT-PCR to examine changes in splice isoform abundance in vehicle-, TEA-, or TTX-treated human iNeurons. sTDP43 isoforms were not only detectable in iNeurons, but also significantly upregulated by treatment with TEA (Figure 3C); while there was a trend towards downregulation with TTX, this did not reach statistical significance. These results suggest that the bidirectional change in N-terminal TDP43 observed in TEA- or TTX-treated iNeurons may be due to altered expression of sTDP43 transcript isoforms. Transcripts encoding flTDP43 were significantly more abundant than the sTDP43 isoforms, and demonstrated a similar upregulation with TEA and trend downwards with TTX (Figure 3C,D). Thus, although all *TARDBP* transcript variants increase with neuronal activity, only sTDP43 isoforms demonstrate corresponding changes at the protein level, perhaps due to selective autoregulation or nuclear retention of spliced full-length transcripts (61–63).

**Figure 3.**
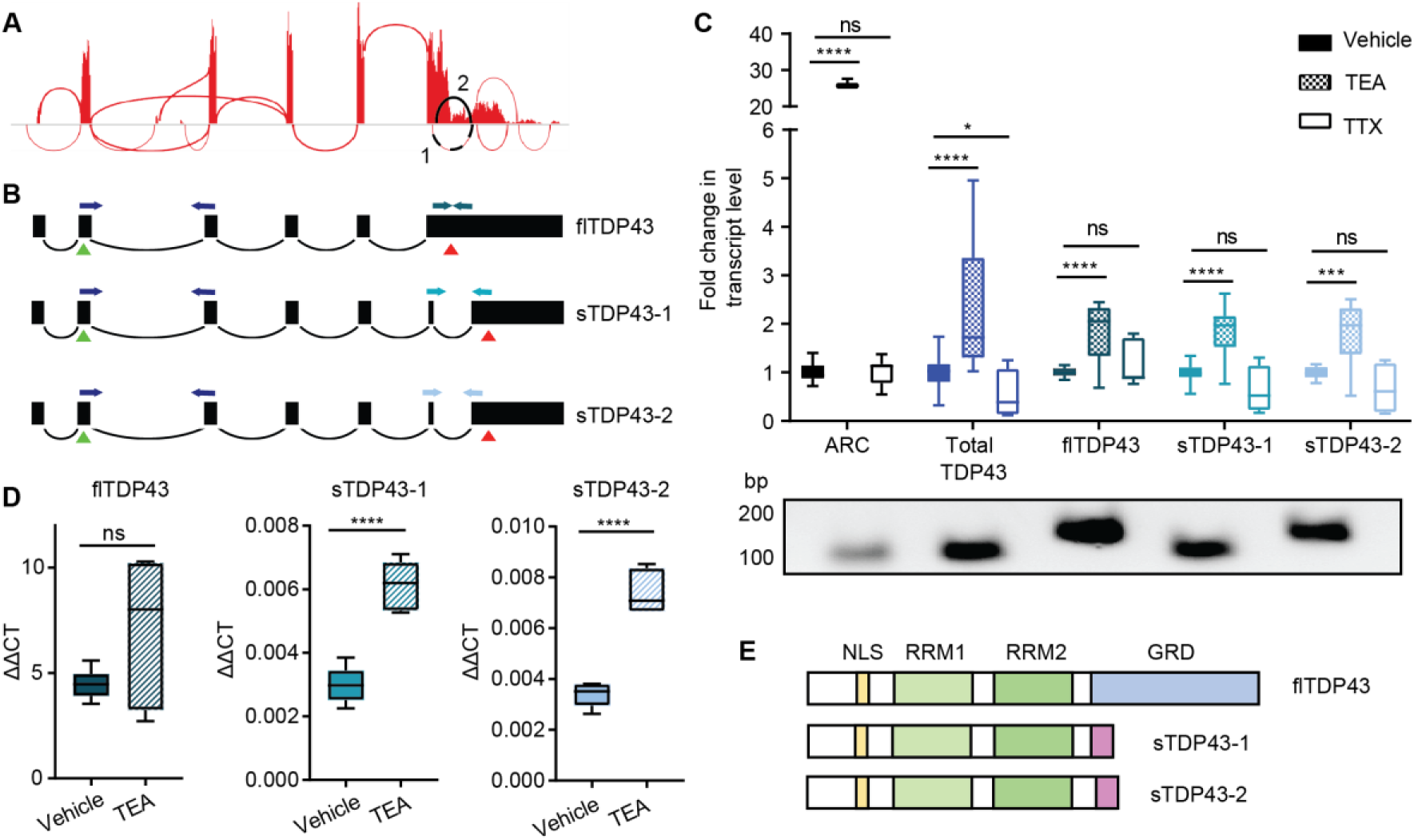
Hyperactivity drives alternative splicing of *TARDBP*. (**A**) Sashimi plot depicting splicing events for the *TARDBP* gene, assembled from HEK293T cell RNA-seq data. Splicing events predicted to skip the majority of exon 6—encoding the TDP43 C-terminus—are highlighted in black. (**B**) Schematic of transcripts predicted to result in full-length (fl) TDP43 and C-terminally shortened (s) TDP43. Green triangles indicate start codons, red triangles indicate stop codons, and PCR primers are color-coded. (**C**) qRT-PCR of human iNeurons treated with TEA or TTX, showing activity-dependent upregulation of total and sTDP43 or downregulation of sTDP43, respectively. ARC (activity related cytoskeleton associated protein) acts as a positive control for activity-dependent gene regulation. PCR products corresponding to each primer pair are shown below. Data combined from 3 replicates, *p<0.05, ***p<0.001, ****p<0.0001, one-way ANOVA with Dunnett’s post-test. (**D**) Raw ΔΔCT values demonstrate the relatively low abundance of sTDP43-1 and −2 transcripts in iNeurons compared to flTDP43. (**E**) Schematic comparing flTDP43 and sTDP43 proteins. The new sTDP43 C-terminus is shown in purple; NLS, nuclear localization signal; RRM, RNA-recognition motif; GRD, glycine rich domain.

The two sTDP43 transcripts (sTDP43-1 and −2) encode proteins that differ by only 3 amino acids, and both lack residues that correspond to the entirety of the glycine rich domain (residues 281-414 of flTDP43) (64). Usage of the common splice acceptor for sTDP43-1 and −2 located within the *TARDBP* 3’UTR results in the inclusion of a new exon encoding a unique 18-amino acid C-terminus not found in flTDP43 (Figure 3E). These splicing events and the distinct sTDP43 C-terminus are highly conserved at both the transcript (Supplemental Table 1) and protein (Supplemental Table 2) levels in humans, non-human primates, and lesser mammals. Despite this, and the previous identification of sTDP43 splice variants in human and murine tissues (60, 61, 64, 65), the pathways governing their expression remain unknown. Our results demonstrate that these variants are dynamically and bidirectionally regulated by neuronal activity, with neuronal hyperactivity resulting in a significant upregulation of sTDP43 at the RNA and protein levels.

### sTDP43 is cytoplasmically localized due to a putative NES in its C-terminal tail

To investigate sTDP43 localization, we transfected rodent primary mixed cortical neurons with diffusely localized mApple to enable visualization of neuronal cell bodies and processes, as well as constructs encoding flTDP43 or sTDP43-1 isoforms fused to an EGFP tag. We then imaged cultures by fluorescence microscopy to examine the localization of each isoform. flTDP43 appeared to be primarily nuclear in distribution, as expected, but sTDP43 demonstrated prominent cytoplasmic deposition (Figure 4A). The dramatic difference in sTDP43 localization was unanticipated given the presence of an intact nuclear localization signal (NLS) within the sTDP43 N-terminus (Figure 3E), and hinted at the presence of a potential nuclear export signal (NES) within the new sTDP43 C-terminus.

**Figure 4.**
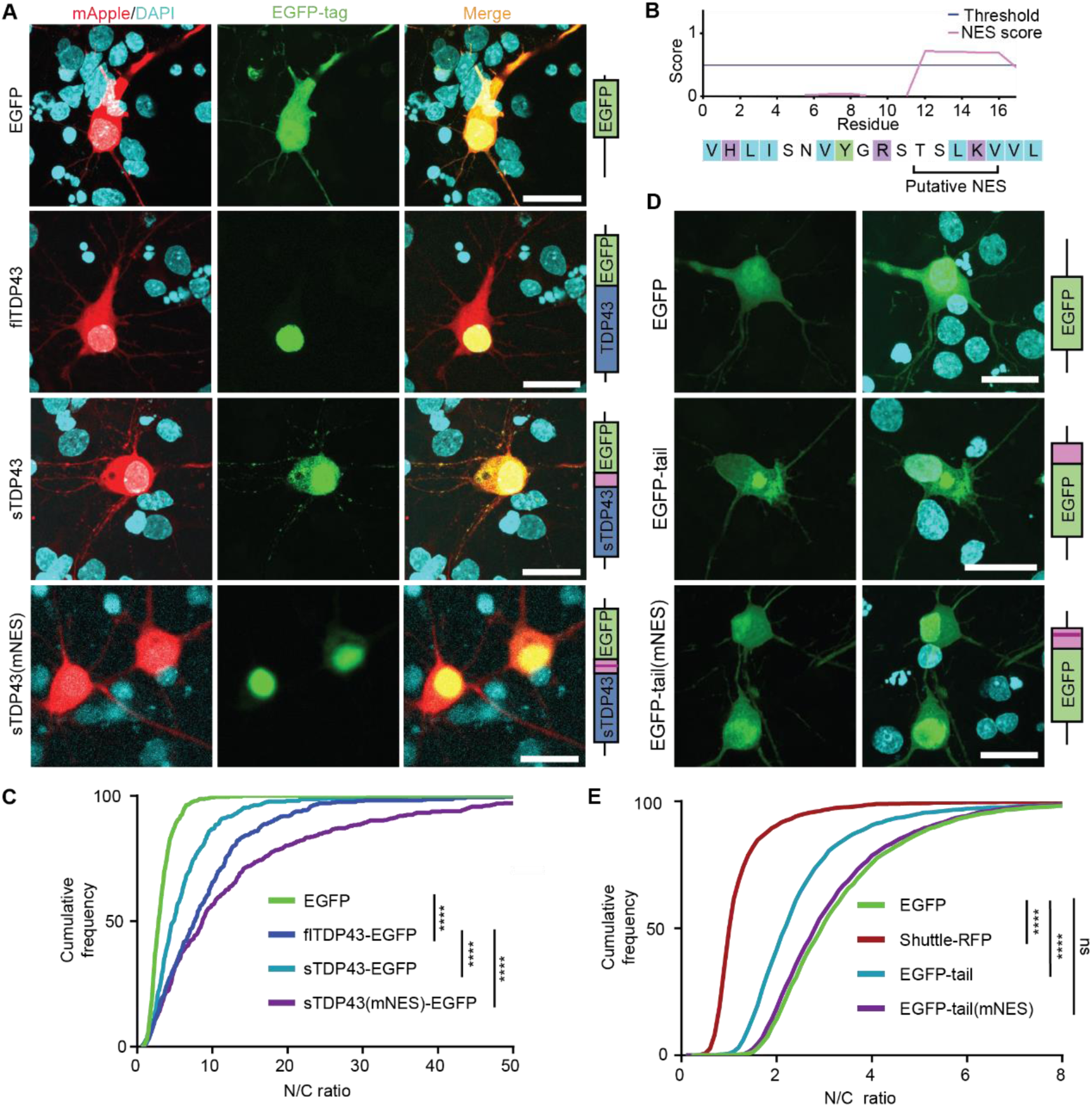
sTDP43 accumulates within the cytoplasm due to a putative NES. (**A**) Rodent primary mixed cortical neurons were transfected with mApple and EGFP-tagged TDP43 isoforms, then imaged by fluorescence microscopy. (**B**) Amino acid sequence of the sTDP43 tail showing the putative NES identified using NetNES 1.1. Light blue, polar; purple, positively charged; green, hydrophobic residues. (**C**) sTDP43-EGFP was significantly more cytoplasmic compared to flTDP43-EGFP, while mutation of the putative NES (mNES) restores nuclear localization. EGFP n=481, flTDP43-EGFP n=385, sTDP43-EGFP n=456, sTDP43(mNES)-EGFP n=490, stratified among 3 replicates, ****p<0.0001, one-way ANOVA with Dunnett’s post-test. (**D**) Rodent primary mixed cortical neurons were transfected with one of three constructs: EGFP alone, EGFP fused to the sTDP43 C-terminus, or EGFP fused to the sTDP43 C-terminal tail harboring a mutated NES (mNES). (**E**) The sTDP43 C-terminus mislocalizes EGFP to the cytoplasm, and mislocalization depends on the putative NES. Shuttle-RFP, a construct with a strong NES, serves as a positive control for a cytoplasmic protein. EGFP n=2490, Shuttle-RFP n=2073, EGFP-tail n=1956, EGFP-tail(mNES) n=2482, stratified among 3 replicates, ****p<0.0001, one-way ANOVA with Dunnett’s post-test. Scale bars in (**A**) and (**D**), 20 µm.

To explore this possibility, we utilized NetNES1.1, an algorithm that employs neural networks and hidden Markov models to predict NES-like motifs from protein primary structure (66). This analysis uncovered a series of amino acids near the sTDP43 C-terminal pole that could potentially act as an NES (Figure 4B). We then tested the function of this putative NES through three complementary experiments. First, we altered the putative NES within sTDP43 by site-directed mutagenesis (TSLKV→GGGGG) and expressed this construct (sTDP43(mNES)) in rodent primary neurons (Figure 4A). Protein localization was assessed by automated microscopy, using scripts that measure fluorescence separately within cytoplasmic and nuclear ROIs, and calculate a nuclear-cytoplasmic ratio (NCR) for TDP43 in each transfected neuron (14, 15). While sTDP43 was localized to both the nucleus and cytoplasm, sTDP43(mNES) displayed a primarily nuclear distribution, more so even than flTDP43, suggesting that the putative NES is necessary for cytoplasmic deposition of sTDP43 (Figure 4C). Second, we fused EGFP to the 18-amino acid tail of sTDP43 (EGFP-tail), or a version of the sTDP43 tail harboring a mutated NES (EGFP-tail(mNES)) (Figure 4D). For comparison, we also expressed Shuttle-RFP, a construct with a strong NES and a weak NLS that exhibits a predominant cytoplasmic distribution (67). Addition of the sTDP43 tail was sufficient to partially exclude EGFP-tail from the nucleus, but this change in distribution was eliminated by mutating the residues making up the putative NES in EGFP-tail(mNES) (Figure 4E). Lastly, we asked whether sTDP43’s cytoplasmic distribution arises from the absence of a nuclear retention signal encoded within the canonical TDP43 C-terminus (68), or the presence of an active NES within the sTDP43 tail. Fusing the sTDP43 tail to flTDP43 markedly shifted the distribution of flTDP43 to the cytoplasm (Supplemental Figure 3), suggesting that sTDP43 localization is dictated primarily by the C-terminal NES, and not the lack of a nuclear retention signal. Together, these data indicate that the distinct sTDP43 C-terminus encodes a functional NES that facilitates cytoplasmic accumulation of sTDP43.

### sTDP43 overexpression is neurotoxic

TDP43 mislocalization is a widely observed phenomenon in ALS, and cytoplasmic TDP43 is a strong predictor of cell death (14). Given these data and the largely cytoplasmic localization of sTDP43, we surmised that sTDP43 accumulation would be toxic to mammalian neurons. We therefore utilized automated microscopy in conjunction with survival analysis to track individual neurons prospectively over time and determine their risk of death in an unbiased and high-throughput manner (14, 15, 59, 69, 70). Rodent primary mixed cortical neurons were transfected with mApple and EGFP-tagged TDP43 isoforms and imaged by fluorescence microscopy at 24h intervals for 10d (71). Custom scripts were used to automatically generate ROIs corresponding to each cell and determine time of death based on rounding of the soma, retraction of neurites, or loss of fluorescence (Figure 5A). The time of death for individual neurons was used to calculate the risk of death in each population relative to a reference group, in this case neurons expressing EGFP (71, 72). In keeping with the results of previous studies, flTDP43 overexpression resulted in a significant increase in the risk of death in comparison to EGFP alone (p<2×10^-16^). sTDP43-1 overexpression elicited an analogous increase in the risk of death for transfected neurons (p<2×10^-16^), suggesting that sTDP43 and flTDP43 display similar toxicity when overexpressed in neurons (Figure 5B).

**Figure 5.**
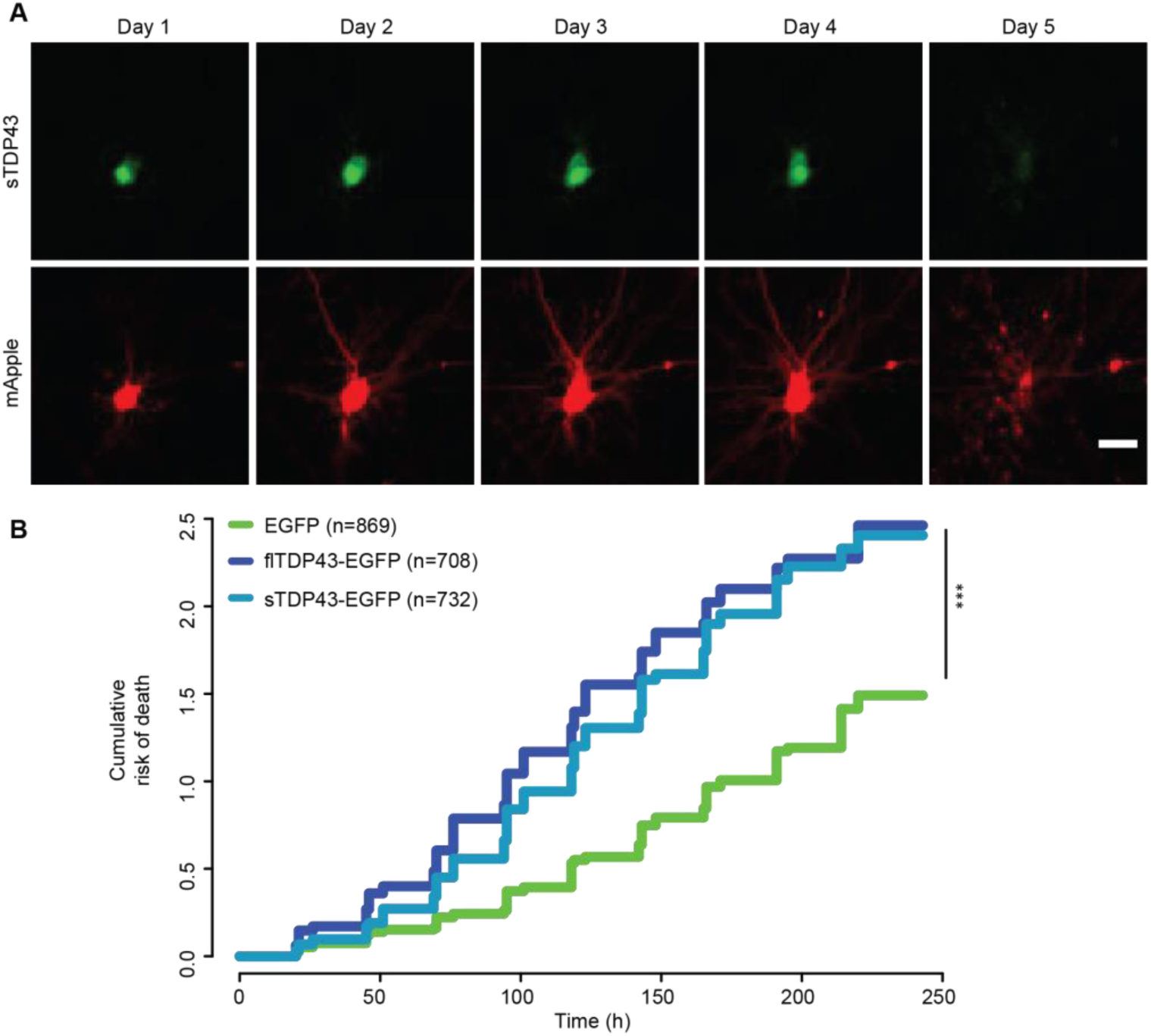
sTDP43 overexpression is neurotoxic. (**A**) Example of a single neuron expressing mApple and sTDP43-EGFP, tracked by longitudinal fluorescence microscopy. Fragmentation of the cell body and loss of fluorescence indicates cell death. (**B**) The risk of death was significantly greater in neurons overexpressing sTDP43-EGFP and flTDP43-EGFP, compared to those expressing EGFP alone. EGFP n= 869, flTDP43-EGFP n=708, sTDP43-EGFP n=732, stratified among 3 replicates, ***p<2×10^-16^, Cox proportional hazards analysis. Scale bar in (**A**), 20 µm.

### sTDP43 alters endogenous TDP43 localization

TDP43 dimerizes via its N-terminus (52, 73–79), and because sTDP43 is exported from the nucleus and contains an intact N-terminus we questioned whether sTDP43 might bind to and sequester endogenous flTDP43 within the cytoplasm. To determine if sTDP43 is capable of interacting with endogenous flTDP43, we transfected HEK293T cells with HaloTag-labeled sTDP43-1 or flTDP43 and isolated the fusion proteins using HaloLink resin (Figure 6A). We detected equivalent amounts of endogenous flTDP43 in eluates from sTDP43-HaloTag and flTDP43-HaloTag, indicating that sTDP43 effectively binds endogenous flTDP43 (Figure 6B).

**Figure 6.**
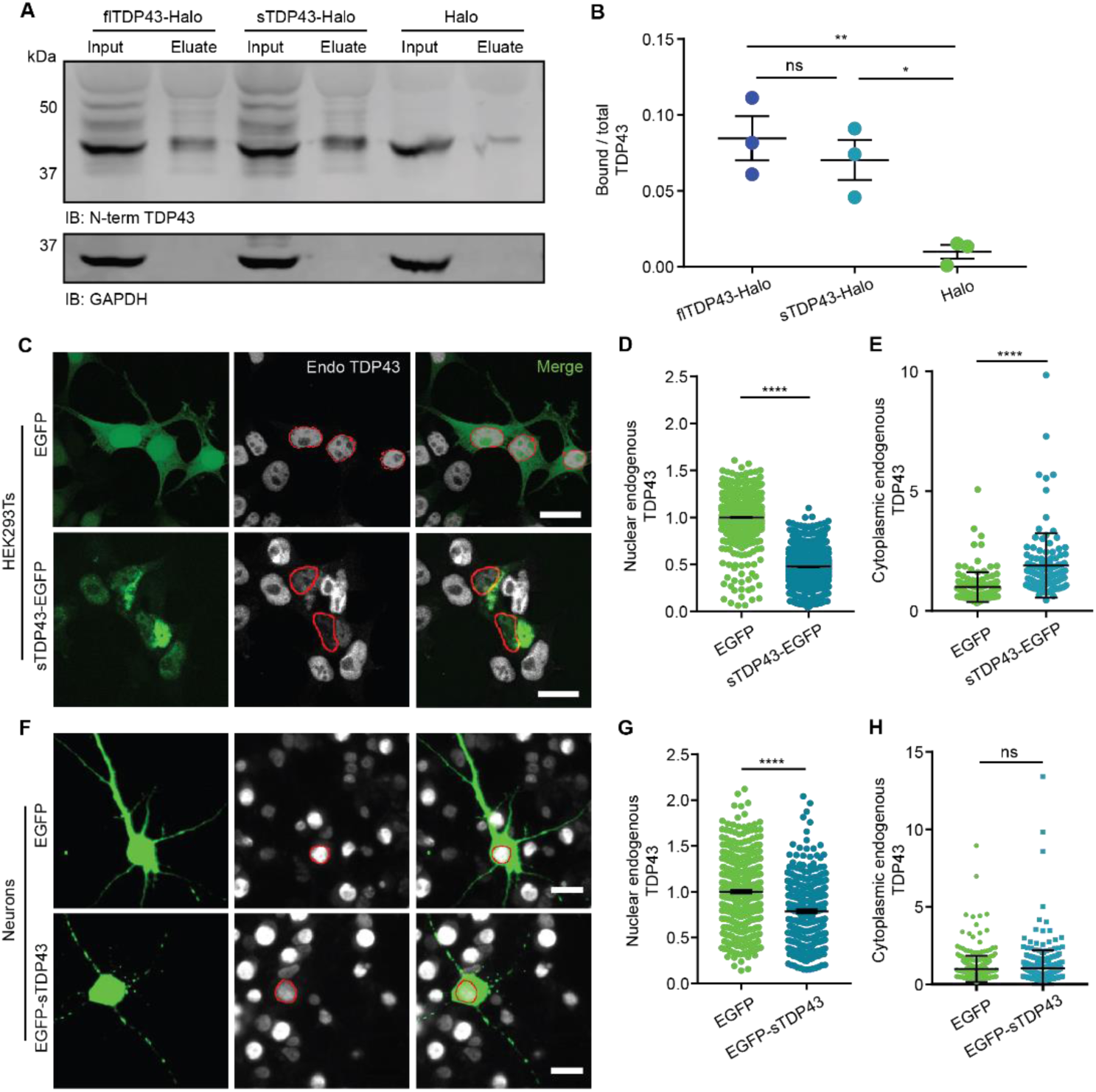
sTDP43 overexpression leads to the cytoplasmic deposition and nuclear clearance of endogenous TDP43. (**A**) HaloTag fusions of flTDP43 or sTDP43 were expressed in HEK293T cells and immunoprecipitated with HaloLink. Bound TDP43 was immunoblotted with an N-terminal TDP43 antibody. GAPDH served as a loading control. (**B**) Quantification of data shown in (A), demonstrating the fraction of total TDP43 bound to flTDP43-Halo, sTDP43-Halo, or Halo alone. Data was combined from 3 replicates, *p<0.05, **p<0.01, one-way ANOVA with Dunnett’s post-test. (**C**) HEK293T cells were transfected with EGFP or EGFP-tagged sTDP43, then immunostained using an antibody that recognizes the endogenous TDP43 C-terminus (Endo). Red, nuclear regions of interest (ROIs) determined by DAPI staining. (**D**) Nuclear, endogenous TDP43 is reduced by sTDP43 overexpression in HEK293T cells. EGFP n=1537, sTDP43-EGFP n=1997, 3 replicates, ****p<0.0001, two-tailed t-test. (**E**) Cytoplasmic endogenous TDP43 is elevated by sTDP43 overexpression in HEK293T cells. EGFP n=129, sTDP43-EGFP n=113, 3 replicates, ****p<0.0001, two-tailed t-test. (**F**) Primary mixed rodent cortical neurons were transfected with EGFP or EGFP-tagged sTDP43, then immunostained using a C-terminal TDP43 antibody. Red, nuclear ROIs determined by DAPI staining. (**G**) sTDP43 overexpression resulted in a significant drop in nuclear, endogenous TDP43 in primary neurons (EGFP n=395, EGFP-sTDP43 n=323, 3 replicates, ****p<0.0001, two-tailed t-test), but this was not accompanied by increases in cytoplasmic, endogenous TDP43 (**H**) (EGFP=394, EGFP-sTDP43=323, 3 replicates, ns by two-tailed t-test). Scale bar in (**C**), (**F**) 20 µm.

We also examined the interaction between sTDP43 and endogenous TDP43 by ICC. HEK293T cells were transfected with EGFP or EGFP-tagged sTDP43, immunostained using a C-terminal TDP43 antibody that recognizes endogenous TDP43 but not sTDP43, and imaged by confocal fluorescence microscopy (Figure 6C). HEK293T cells overexpressing EGFP-tagged sTDP43 displayed cytoplasmic inclusions that strongly colocalize with endogenous TDP43. Moreover, we observed significant reductions in nuclear endogenous TDP43 in association with cytoplasmic TDP43 deposition (Figure 6D, E), suggesting cytoplasmic sequestration of endogenous TDP43 by sTDP43. Rodent primary mixed cortical neurons overexpressing sTDP43-1 displayed a similar depletion of endogenous TDP43 from the nucleus (Figure 6F, G). Thus, sTDP43 overexpression results in both cytoplasmic deposition and nuclear clearance of endogenous TDP43, recapitulating signature features of ALS pathology and implying that both gain- and loss-of-function mechanisms contribute to toxicity.

In sTDP43-transfected cells, we observed significant variability in the degree of TDP43 nuclear exclusion and cytoplasmic aggregation, which we suspected was due to differences in sTDP43 expression among transfected cells. Because the abundance of a fluorescently-labeled protein is directly proportional to the intensity of the fluorescent tag (80), we estimated sTDP43 expression in each cell by measuring single-cell EGFP intensity, and separated cells into 5 bins based on sTDP43 expression level. In doing so, we observed a direct relationship between the extent of endogenous TDP43 mislocalization and sTDP43 expression (Supplemental Figure 4). These results indicate that TDP43 pathology may become more prevalent as sTDP43 expression is upregulated by neuronal hyperexcitability or other mechanisms.

We failed to observe significant increases in cytoplasmic TDP43 deposition in transfected primary neurons (Figure 6H), potentially due to steric inhibition of sTDP43 localization and function by fusion with EGFP or HaloTag (Supplemental Figure 5). Placing the EGFP tag on the C-terminus of sTDP43 partially prevented cytoplasmic localization of sTDP43-EGFP (Supplemental Figure 5A-C), likely by masking the C-terminal NES. Similarly, we found that fusion of HaloTag with the N-terminus of sTDP43 significantly inhibits its binding to endogenous TDP43 (Supplemental Figure 5D, E). As such, N-terminal labeling of sTDP43 leaves the NES accessible but blocks association with endogenous TDP43, while C-terminal sTDP43 labeling obstructs the NES but allows interaction with endogenous TDP43.

### sTDP43 lacks canonical functions of flTDP43

To further examine the possibility that sTDP43 elicits loss-of-function toxicity, we assessed the ability of sTDP43 to participate in TDP43-related splicing activity. In keeping with TDP43’s function as a splicing repressor, TDP43 effectively blocks the inclusion of cystic fibrosis transmembrane conductance regulator (CFTR) exon 9 (81, 82) (Supplemental Figure 6A). In HEK293T cells expressing a CFTR minigene reporter, cotransfection with EGFP-flTDP43 resulted in proficient exon 9 exclusion as measured by PCR. In contrast, EGFP-sTDP43-1 expression failed to significantly affect exon 9 exclusion (Supplemental Figure 6B,C), suggesting that without the C-terminus, sTDP43 is incapable of TDP43-specific splicing (60, 64, 65).

Functional flTDP43 also participates in an autoregulatory feedback loop, in which flTDP43 recognizes sequences within the *TARDBP* 3’UTR, triggering alternative splicing and/or polyadenylation and subsequent mRNA degradation (83, 84). To determine if sTDP43 is able to regulate endogenous TDP43 expression via this mechanism, we employed a TDP43 autoregulation reporter consisting of an open reading frame (ORF) encoding the fluorescent protein mCherry upstream of the *TARDBP* 3’ UTR (69) (Supplemental Figure 7A). In rodent primary cortical neurons expressing the TDP43 autoregulation reporter, cotransfection with EGFP-tagged flTDP43 resulted in a decrease in reporter signal, as expected. EGFP-labeled sTDP43-1 displayed lesser effects on reporter fluorescence, suggesting that its ability to autoregulate TDP43 is impaired (Supplemental Figure 7B, C). Likewise, when expressed in HEK293T cells, sTDP43-1 exhibited a similarly muted effect on endogenous TDP43 at the transcript and protein level (Supplemental Figure 7D-F), consistent with poor autoregulation. Together, these results indicate that sTDP43 lacks many of the canonical functions of TDP43, including its splicing and autoregulatory abilities.

### sTDP43 colocalizes with markers of stress granules

Previous studies suggested that sTDP43 associates with protein components of cytoplasmic stress granules, including G3BP1 and TIA1 (65). Therefore, we immunostained for G3BP1 and TIA1 in HEK293T cells overexpressing EGFP-tagged sTDP43-1, before and after application of osmotic stress (0.4M sorbitol). Prior to sorbitol treatment, we noted substantial colocalization of sTDP43-1 with G3BP1 (Supplemental Figure 8A) and TIA1 (data not shown) in large cytoplasmic deposits; these structures were unique to cells transfected with sTDP43-1, suggesting that sTDP43 overexpression elicits the formation of irregular structures rich in stress granule components. However, when cells were stressed with 0.4M sorbitol we observed the formation of multiple small, punctate structures resembling stress granules that colocalize with both G3BP1 and TIA1, as well as endogenous TDP43 (Supplemental Figure 8B). Moreover, while osmotic stress drives flTDP43 to the cytoplasm, it has little effect on sTDP43-1 localization (Supplemental Figure 8C). These data confirm that sTDP43 localizes to stress granules, and further imply that sTDP43 production may be sufficient for the assembly of cytoplasmic stress granule-like structures even in the absence of stress.

### sTDP43 transcripts are enriched in murine and human lumbar motor neurons

To determine if sTDP43 isoforms are produced *in vivo* and assess their expression in different regions of the central nervous system, we took advantage of a previous study that analyzed the transcriptome from murine frontal cortex and lumbar spinal motor neurons isolated by laser capture microdissection (Figure 7A) (85). The most abundant splice isoform in frontal cortex homogenate was flTDP43, with predominant use of the conventional termination codon within *TARDBP* exon 6. However, splicing of the *TARDBP* locus, and in particular exon 6 and the 3’UTR, was dramatically altered in murine spinal motor neurons. In contrast to what was observed in frontal cortex, two splicing events corresponding to sTDP43 variants 1 and 2 were strongly favored in spinal motor neurons—these isoforms were upregulated ∼12- and 10-fold, respectively, in lumbar spinal neurons relative to frontal cortex (Figure 7B, C). Further, while an ALS-associated *TARDBP* mutation (Q331K) did not affect sTDP43 transcript abundance, we noted age-related increases in sTDP43 mRNA levels in 20-month vs. 5-month old mouse cortices (Supplemental Figure 9). These data show that sTDP43 isoforms are not only detectable *in vivo* within the murine CNS, but they are also significantly upregulated by age and enriched in spinal motor neurons, a cell type selectively targeted in ALS.

**Figure 7.**
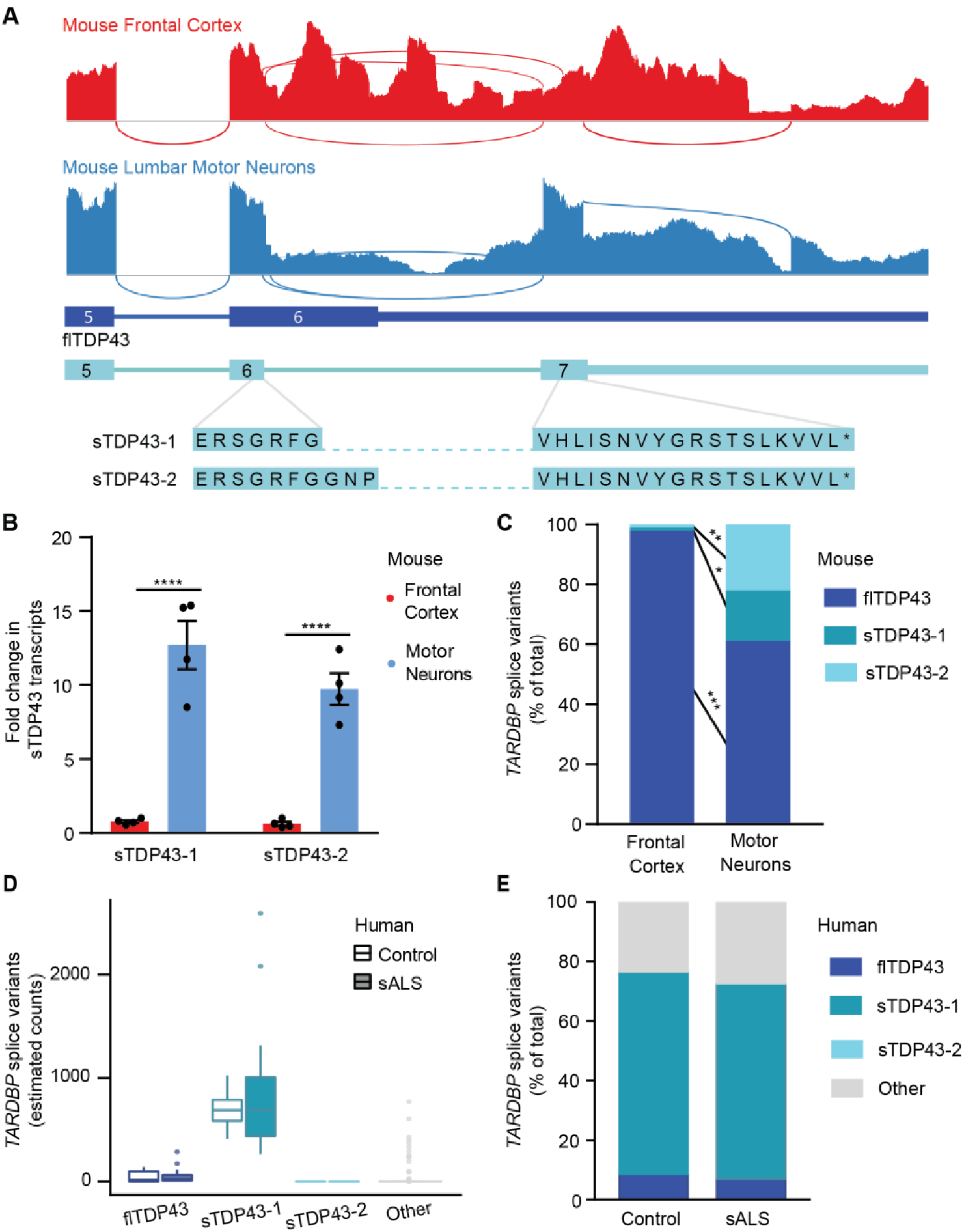
sTDP43 transcripts are enriched in lumbar motor neurons. (**A**) Sashimi plots depicting *TARDBP* splicing in murine frontal cortex homogenate (red) or microdissected lumbar motor neurons (blue). (**B**) Both sTDP43-1 and sTDP43-2 splice events are highly enriched in lumbar motor neurons compared to frontal cortex homogenate. Graph depicts read counts normalized to reads per million for each library. Analysis includes 4 replicates, ****p<0.0001 multiple t-test with the Holm-Sidak correction. (**C**) While sTDP43-1 and −2 each comprise ∼1% of the total *TARDBP* transcripts in frontal cortex homogenate, they make up 17 and 22% of total *TARDBP* transcripts in lumbar motor neurons, respectively. Frontal cortex n=6, lumbar motor neurons n=4, *p<0.05, **p<0.01, ***p<0.001, two-way ANOVA with Sidak’s multiple comparison test. (**D**) sTDP43-1 is enriched within lumbar motor neurons microdissected from both control (n=9) and sALS (n=13) patient tissue. (**E**) sTDP43-1 makes up the majority of total *TARDBP* transcripts in both control and sALS patient lumbar motor neurons.

We also examined sTDP43 expression in human spinal neurons utilizing published RNA-seq data from laser-captured lumbar spinal motor neurons, isolated from control and sporadic (s) ALS patient tissue (86). Within this dataset, we identified specific transcripts corresponding to flTDP43, sTDP43-1, and sTDP43-2, and characterized the remaining *TARDBP* variants as “other.” Although there was no apparent difference in the abundance of any *TARDBP* transcripts between sALS and control motor neurons, we noted a dramatic enrichment of sTDP43-1 transcripts in human spinal neurons, in comparison to flTDP43, sTDP43-2, and other *TARDBP* variants (Figure 7D, E). Furthermore, and in contrast to what we observed in rodent spinal neurons, sTDP43-1 was the predominant *TARDBP* splice isoform detected in human spinal neurons. To extend these findings, we also examined available RNA-seq data from spinal cord ventral horn homogenate from control and sALS patients (87), as well as cerebellum and frontal cortex from controls, individuals with sALS, and patients bearing disease-associated mutations in *C9ORF72* (C9ALS) (88). Despite fundamental differences in sample preparation and sequencing methodology that prevented the direct comparison of transcript abundance between tissue types (Supplemental Figure 10A), we consistently observed significant expression of sTDP43-1 but not sTDP43-2 in several different regions of the CNS, including but not limited to spinal motor neurons, cerebellum, and frontal cortex, though there were no differences between control, sALS or C9ALS patient samples (Supplemental Figure 10B, C).

### Endogenously produced sTDP43 is detectable using specific antibodies

To distinguish natively-produced sTDP43 species, we generated an antibody directed against the unique 18-amino acid C-terminus of sTDP43 (Figure 3E). This antibody specifically recognized EGFP fused to the sTDP43 C-terminus, suggesting that the sTDP43 tail is sufficient for immunoreactivity, and the signal was completely abolished by preincubation with the immunizing peptide (Supplemental Figure 11A). Furthermore, expression of artificial miRNAs (amiRNAs) targeting TDP43 (69, 89) effectively reduced flTDP43 levels, as expected (Supplemental Figure 11B, C), and also decreased sTDP43 immunoreactivity (Supplemental Figure 11D, E), confirming antibody specificity. We further validated the sTDP43 antibody by transfecting HEK293T cells with EGFP-tagged sTDP43-1, isolating RIPA- and urea-soluble protein fractions, and immunoblotting for sTDP43. In previous studies, overexpressed sTDP43 was highly insoluble (64); supporting this, we detected EGFP-sTDP43 exclusively in the urea-soluble fraction while EGFP-flTDP43 appeared in both RIPA- and urea-soluble fractions (Figure 8A). We also tested the sTDP43 antibody in human iNeurons treated with TEA or TTX to induce or abolish neuronal activity, respectively (Figure 8B). In these studies, TEA increased sTDP43 immunoreactivity, while TTX reduced sTDP43 levels (Figure 8C), consistent with activity-dependent upregulation of N-terminally labeled D2-TDP43 (Figure 2) and its detection by antibodies specific for the TDP43 N-terminus (Figure 1). Notably, sTDP43 antibodies detected numerous cytoplasmic puncta in TEA-treated neurons that were less apparent in vehicle- and TTX-treated cells, and the background nuclear signal was minimal in all cases. Identical sTDP43-positive cytoplasmic puncta were observed in rodent primary mixed cortical neurons treated with bicuculline (Supplemental Figure 11F). These data indicate that sTDP43 antibodies selectively detect truncated, cytoplasmic, and insoluble TDP43 species by western blot and ICC, establishing them as useful tools for investigating sTDP43 deposition and its potential role in neurodegeneration.

**Figure 8.**
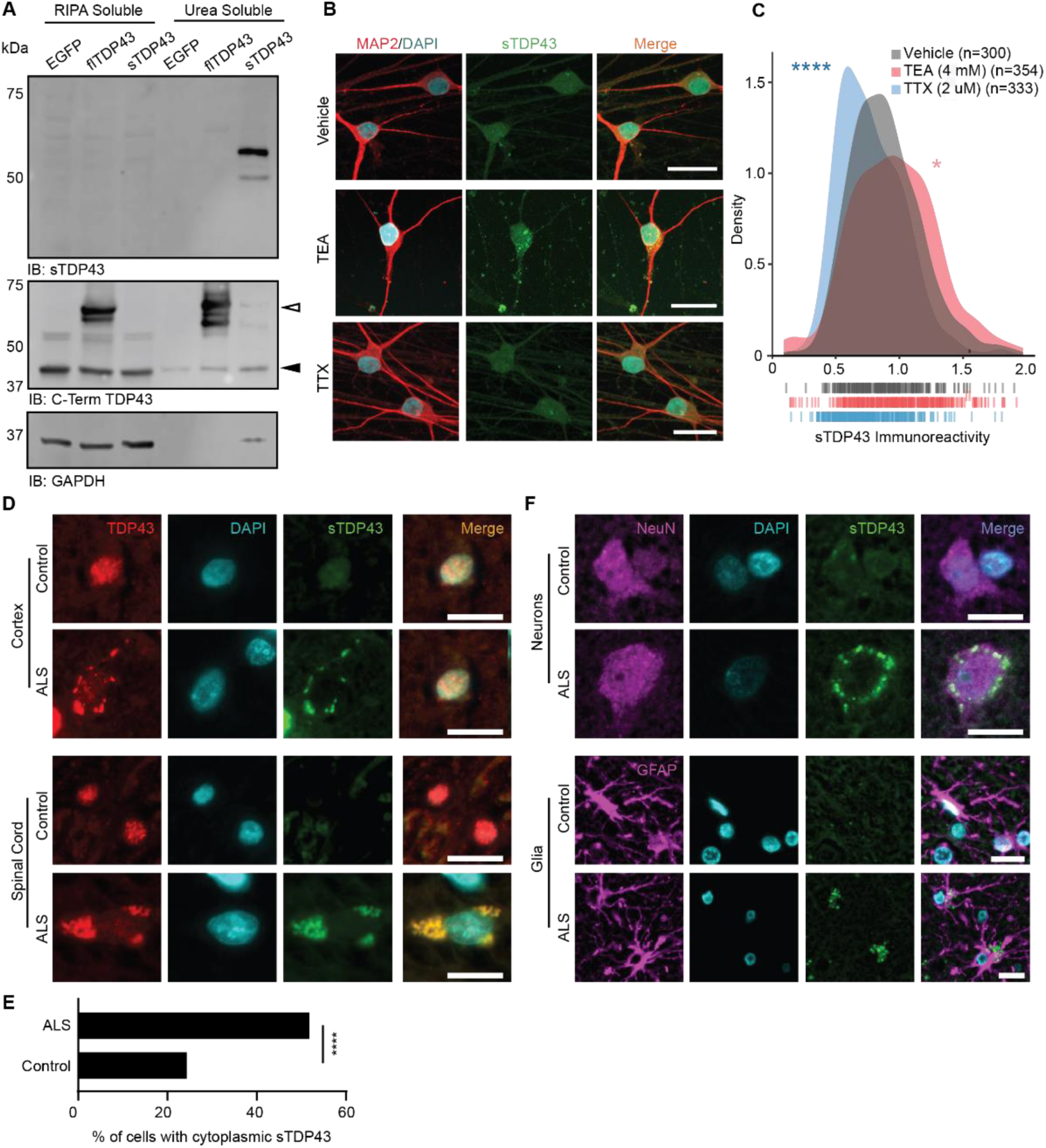
Endogenous sTDP43 is detectable *in vivo* by antibodies generated against its distinct C-terminus. (**A**) Western blot of EGFP-tagged flTDP43 or sTDP43 overexpressed in HEK293T cells. Black arrowhead, endogenous TDP43; white arrowhead EGFP-flTDP43. (**B**) ICC using sTDP43 antibodies showed increased immunoreactivity in TEA-treated iNeurons and decreased immunoreactivity in TTX-treated iNeurons. (**C**) Density plot depicting the change in sTDP43 immunoreactivity between conditions. Vehicle n=300, TEA n=354, TTX n=333, 3 replicates, dashes indicate single neurons, *p<0.05, ****p<0.0001, Kolmogorov-Smirnov test. (**D**) IHC comparing the distribution of N-terminal TDP43 and sTDP43 in spinal cord and cortex from patients with sporadic ALS and controls. (**E**) Quantification of cells with cytoplasmic sTDP43 in control and ALS patient spinal cord (control n=115, ALS n=110, data representative of two control and three ALS patients, ****p<0.0001, Fisher’s exact test). (**F**) IHC demonstrating neuronal and glial sTDP43 accumulation in cortex from individuals with sALS and controls. Scale bars in (**B**), (**D**), and (**F**) 20 µm.

Based on the observed upregulation of sTDP43 splice isoforms in lumbar spinal neurons, we employed our newly-developed sTDP43 antibody for detecting sTDP43 *in vivo* within murine spinal cord sections. As predicted from the RNA-sequencing data, we detected cytoplasmic sTDP43 in anterior horn neurons from the lumbar spinal cord (Supplemental Figure 12A), confirming the subcellular distribution of the protein originally noted *in vitro*. We also observed strong colocalization of sTDP43 with GFAP-positive astrocytic projections within the spinal white matter, indicating astrocytic expression of sTDP43 (Supplemental Figure 12B). Subsequent studies confirmed that sTDP43 is endogenously produced by human iPSC-derived astrocytes (Supplemental Figure 13), suggesting that while sTDP43 is enriched within spinal neurons (Figure 7), it is also synthesized by supporting glia.

### sTDP43 pathology is observed in ALS patient tissue

Given that (a) sTDP43 is endogenously produced at relatively high levels in spinal motor neurons, (b) neuronal hyperexcitability is a conserved feature of ALS, and (c) sTDP43 is upregulated by neuronal activity and age, we suspected that sTDP43 may accumulate in individuals with sALS. To address this question, we immunostained human cortex and spinal cord sections from sALS, C9ALS, and unaffected control patients using antibodies that recognize the TDP43 N-terminus or our newly-developed sTDP43 antibodies (Figure 8D). As predicted, immunostaining with N-terminal TDP43 antibodies showed both a reduction in nuclear signal and the appearance of cytoplasmic inclusions selectively in ALS patient tissue. While control tissue exhibited low immunoreactivity for sTDP43 in both the cortex and spinal cord, we observed a striking accumulation of sTDP43 within cytoplasmic deposits in ALS spinal cord and cortex (Figure 8E). sTDP43-positive inclusions closely colocalized with N-terminally reactive cytoplasmic aggregates but not residual nuclear TDP43, suggesting that sTDP43 antibodies specifically label cytoplasmic deposits in ALS tissue. In this limited case study, sTDP43 pathology appeared to be conserved between sALS and C9ALS (Supplemental Figure 14A), hinting at a conserved process. We also observed a tight correlation between conventional TDP43 pathology (nuclear exclusion and cytoplasmic aggregation) and sTDP43 deposition—in ALS samples, neurons displaying TDP43 nuclear exclusion almost always showed cytoplasmic sTDP43 pathology. Additionally, we noted several cells from ALS patients that exhibited sTDP43 deposits despite a normal nuclear TDP43 pattern, perhaps illustrating an early stage of pathology (Supplemental Figure 14B, C). Even so, we detected significant heterogeneity in sTDP43 pathology among ALS cases, indicating the presence of additional, unknown factors that could impact sTDP43 deposition or immunoreactivity.

In light of endogenous sTDP43 detected within mouse spinal cord astrocytes (Supplemental Figure 12B) and human iPSC-derived astrocytes (Supplemental Figure 13), we asked if sTDP43 pathology might also be present within astrocytes. In sections from controls and sALS patients immunostained with sTDP43 antibodies, neurons and glia were identified by co-staining with NeuN and GFAP antibodies, respectively (Figure 8F). Cytoplasmic sTDP43 accumulations were detected in both NeuN-positive neurons and GFAP-positive astrocytes, suggesting that sTDP43 pathology is not limited to neurons. Taken together, these results demonstrate that endogenous sTDP43 accumulates within neurons and glia of individuals with ALS, supporting a potentially pathogenic contribution of sTDP43 isoforms to ALS pathogenesis.

## Discussion

In this study, we show that neuronal hyperactivity leads to the selective upregulation of C-terminally truncated TDP43 isoforms (sTDP43). These isoforms are intrinsically insoluble and accumulate within cytoplasmic aggregates by virtue of a NES present within a unique 18-amino acid C-terminus. sTDP43 also sequesters endogenous TDP43 within cytoplasmic aggregates and induces its clearance from the nucleus, thereby recapitulating signature pathologic changes found in the majority of individuals with ALS and implicating complementary gain- and loss-of-function mechanisms in disease pathogenesis. sTDP43 transcripts are enriched in spinal motor neurons, a cell type that is selectively vulnerable in ALS, and post-mortem samples from individuals with ALS show conspicuous accumulations of sTDP43 within affected neurons and glia. These observations suggest a fundamental link between neuronal hyperexcitability and TDP43 pathology, two conserved features characteristic of both familial and sporadic ALS. Moreover, they raise the possibility that sTDP43 production and/or its accumulation are heretofore-unrecognized contributors to neurodegeneration in ALS.

A series of previous studies demonstrated that alternative *TARDBP* splicing gives rise to truncated TDP43 isoforms lacking the C-terminus that are highly insoluble when overexpressed in heterologous systems (60, 64, 65). Here, we show for the first time that neuronal activity selectively upregulates these truncated isoforms, which we collectively labeled sTDP43, despite a simultaneous increase in mRNA encoding full-length TDP43. This discrepancy may arise from the relative inability of sTDP43 to effectively participate in autoregulation (Supplemental Figure 7), or the presence of unique elements within the flTDP43 3’UTR leading to nuclear mRNA retention and/or destabilization (61–63, 83, 84). As such, the activity-dependent and apparently selective upregulation of sTDP43, together with the widespread neuronal hyperactivity observed in ALS patients, animal models, and human iPSC-derived neurons (26, 28, 34, 35, 42, 43), may be a crucial factor driving sTDP43 deposition in ALS tissue.

In keeping with previous studies (64, 65), overexpressed sTDP43 accumulates in the cytoplasm where it often forms large, insoluble inclusions. The low-complexity domain (LCD) within the TDP43 C-terminus promotes liquid-phase separation and aggregation (90–94). Even so, our observations and those of others (64, 65) show that sTDP43 is relatively insoluble and prone to aggregation, despite lacking the LCD. A growing body of evidence suggests that proteins with complex, folded domains such as the TDP43 RNA recognition motifs (RRMs) are highly susceptible to aggregation (95). Rather than promoting insolubility, the presence of LCDs within these proteins protects against misfolding and aggregation by enabling reversible phase transitions during conditions of supersaturation. Thus, LCDs may permit higher local concentrations of RRM-containing proteins than would otherwise be possible without misfolding and/or aggregation (96). In this regard, the absence of the LCD may be directly responsible for the enhanced aggregation of sTDP43; indeed, several RNA-binding proteins display similar phenotypes upon removal of the LCD, including PUB1, PAB1 and SUP35 (96–99).

Using predictive software, we identified a potential NES located within the 18-amino acid sTDP43 C-terminus, and experimentally confirmed that this segment drives cytoplasmic sTDP43 localization. This NES appears to be dominant over the functional NLS present within the N-terminus of sTDP43, either due to a high affinity for nuclear exporters or because of enhanced accessibility of the NES at the extreme C-terminus of the protein. The previously annotated TDP43 NES (68, 100) exhibits leucine/isoleucine-rich sequences favored by exportin-1 (XPO1), an essential mediator of protein nuclear export (101). Nevertheless, scant experimental evidence suggests that this sequence functions as a true NES. TDP43 and XPO1 do not interact with one another *in vitro* (15, 102), and unbiased proteomics studies have failed to identify TDP43 as an XPO1 cargo protein (103, 104). Further, TDP43 localization is unaffected by XPO1 inhibition or deletion of the putative NES (15). In contrast, the NES uncovered within the sTDP43 C-terminal tail is both necessary and sufficient for sTDP43 nuclear export, suggesting that it is a *bona fide* NES.

sTDP43 lacks the C-terminal glycine rich domain required for splicing activity (105); as such, sTDP43 is incapable of *CFTR* minigene splicing or effectively participating in TDP43 autoregulation, which involves differential splicing of the *TARDBP* 3’UTR (83, 84). The C-terminal glycine rich domain is also required for toxicity upon TDP43 overexpression in yeast (94). Nevertheless, sTDP43 overexpression was still lethal in neurons. We suspect that sTDP43-related toxicity arises from a combination of factors, including (a) the NES within the new C-terminal tail region provoking cytoplasmic sTDP43 deposition; (b) its interaction with endogenous flTDP43 via its N-terminus (52, 73, 106); and (c) the presence of intact RRMs that enable sTDP43 to bind and potentially sequester cytoplasmic mRNAs.

sTDP43 isoforms are highly conserved in humans, non-human primates, and lesser mammals at the transcript and protein levels. Such evolutionary conservation suggests that these isoforms fulfill unknown functions, perhaps involving a compensatory response to chronic neuronal hyperactivity or generalized stress. Intriguingly, sTDP43 transcripts are significantly enriched in murine motor neurons compared to frontal cortex homogenate, their expression increases with age, and sTDP43-1 is the dominant *TARDBP* species in human lumbar motor neurons, raising the possibility that spinal motor neurons accumulate potentially toxic levels of sTDP43 in response to aging and hyperexcitability. Future studies are needed to determine whether native sTDP43 performs an essential function in motor neurons or other cell types, and if sTDP43 contributes to the selective vulnerability of aged motor neurons in ALS (107, 108).

By creating an antibody that recognizes the unique sTDP43 C-terminus, we detected cytoplasmic sTDP43 inclusions selectively within the spinal cord and cortex of ALS patients, including individuals with sALS and C9ALS. In addition, the presence of sTDP43 deposits coincided with nuclear TDP43 exclusion, as predicted by sTDP43 nuclear export and its affinity for flTDP43. Although the aggregation-prone TDP43 C-terminus forms a core component of the cytoplasmic inclusions found in ALS patients (109–117), emerging evidence suggests that N-terminal TDP43 fragments also contribute to ALS pathogenesis. N-terminal TDP43 fragments are observed in ALS patient spinal cord (118, 119), and in keeping with studies of RNA-binding proteins in yeast, the TDP43 RRMs misfold and aggregate *in vitro* without the C-terminal LCD to maintain solubility (65, 95, 97–99, 120–122). Independent of the RRMs, the TDP43 N-terminus enhances TDP43 aggregation and toxicity (65, 79, 120, 121), potentially adding to sTDP43 insolubility and the impact of sTDP43 deposition in affected neurons.

TDP43-positive cytoplasmic inclusions in ALS are not limited to neurons but are also found in astrocytes and oligodendrocytes (123–126). Astrocytes help regulate extracellular glutamate levels, and their dysfunction in ALS may lead to impaired synaptic glutamate buffering in sporadic as well as familial ALS (127–132). In addition to detecting endogenous sTDP43 production in cultured human astrocytes and murine spinal cord, we noted disease-specific astrocyte sTDP43 pathology in sALS patient tissue. Although the effect of sTDP43 accumulation in these cells remains to be determined, it is possible that sTDP43-induced astrocyte toxicity triggers a feed-forward mechanism in which reduced glutamate buffering results in neuronal hyperactivity, increased sTDP43 production, and subsequent neurodegeneration.

We and others (61) have been unable to reliably detect endogenous sTDP43 isoforms by western blotting, an effect likely related to the intrinsic insolubility of the protein (61, 65) (Figure 8A). In previous studies, however, tandem mass spectroscopy identified C-terminal sequences unique to sTDP43 in mouse brain tissue (61), confirming the presence of sTDP43 in the CNS. Despite the limitations of the newly-developed sTDP43 antibody for western blotting, its utility for immunofluorescence is supported by several observations. First, sTDP43 immunoreactivity varied bidirectionally in TEA- and TTX-treated iNeurons (Figure 8B, C), analogous to the pattern detected by N-terminal but not C-terminal TDP43 antibodies (Figure 1H-K). Second, endogenous TDP43 labeled with Dendra2 at the N-terminus but not C-terminus demonstrated an identical pattern (Figure 2), and nearly all of the sTDP43 immunoreactivity in human brains overlapped with signal from an N-terminally directed TDP43 antibody (Figure 8D), as predicted for sTDP43 isoforms missing the conventional C-terminus.

Our work underlines the significance of previously identified sTDP43 isoforms and highlights a pivotal connection between neuronal hyperexcitability and TDP43 pathology, two conserved findings in ALS. Many questions remain, including the function of sTDP43 isoforms, the extent and pervasiveness of sTDP43 pathology in ALS, and whether cell type- or species-specific differences in sTDP43 expression contribute to the selective vulnerability of human motor neurons in ALS. Complementary investigations of sTDP43 splicing and its regulation are crucial if we are to determine if targeted manipulation of sTDP43 has the potential to prevent or slow motor neuron degeneration in ALS.

## Materials and Methods

Materials and methods are listed in the supplemental information.

### Ethics statement

All vertebrate animal work was approved by the Committee on the Use and Care of Animals (UCUCA) at the University of Michigan and in accordance with the United Kingdom Animals Act (1986). All experiments were performed in accordance with UCUCA guidelines. Rats (*Rattus norvegicus*) used for primary neuron collection were housed singly in chambers equipped with environmental enrichment. All studies were designed to minimize animal use. Rats were cared for by the Unit for Laboratory Animal Medicine at the University of Michigan; all individuals were trained and approved in the care and long-term maintenance of rodent colonies, in accordance with the NIH-supported Guide for the Care and Use of Laboratory Animals. All personnel handling the rats and administering euthanasia were properly trained in accordance with the UM Policy for Education and Training of Animal Care and Use Personnel. Euthanasia was fully consistent with the recommendations of the Guidelines on Euthanasia of the American Veterinary Medical Association.

## Supporting information

Weskamp et al. 2019 Supplemental Information

## Acknowledgements

This work was supported by National Institutes of Health (NIH), National Institute for Neurological Disorders and Stroke (NINDS) R01-NS097542, National Institute for Aging (NIA) P30 AG053760 (SJB), the University of Michigan Protein Folding Disease Initiative, and Ann Arbor Active Against ALS. We thank Dr. M. Uhler for advice, protocols and reagents; Drs. M. Ward, Z. Xu, and Y. Ayala for reagents; and Dr. J. Parent, Mr. M. Malik, Ms. T. Garay, and Mr. M. McMillian for their suggestions. We also thank Mr. M. Perkins from the Michigan Brain Bank (5P30 AG053760, University of Michigan Alzheimer’s Disease Core Center), the Michigan ALS Biorepository, the University of Michigan DNA Sequencing Core, and the University of Michigan Department of Pharmacology for access to their confocal microscopy core. Finally, we thank those that donated the fibroblast and tissue samples that made these studies possible.

## Author Contributions

K.W. was responsible for conceptualization, methodology, investigation, formal analysis, writing and visualization. S.J.B. contributed to conceptualization, methodology, formal analysis, writing, visualization, supervision, project administration, and funding acquisition. E.M.T. contributed to conceptualization and methodology, and R.M. was responsible for software. M.A.W. and N.B.G. contributed to data curation and formal analysis. J.S. and A.S. contributed to supervision and project administration. J.P.M., Z.L., C.M.G., and A.S. contributed to investigation.

