## Supplementary material for "Shortened TDP43 isoforms upregulated by neuronal hyperactivity drive TDP43 pathology in ALS": Weskamp et al. 2019 Supplemental Information

*** To whom correspondence should be addressed:**

Sami Barmada

### Supplemental Materials and Methods

#### Generation and maintenance of iPSCs

Fibroblasts were reprogrammed into iPSCs via transfection with episomal vectors encoding seven reprogramming factors (1) and validated as previously described (2) (Supplemental Table 7). All iPSC lines were cultured in Essential 8 (E8) media (Gibco A1517001) on plates coated with vitronectin (Gibco A14700) diluted 1:100 in $Mg^{2+}$/$Ca^{2+}$-free phosphate buffered saline (PBS, Gibco 14190-144). Cells were passaged every 5–6d using 0.5 mM EDTA (Sigma E7889) dissolved in PBS followed by gentle trituration in E8 media with a P1000 pipette. All lines are verified mycoplasma-free on a monthly basis.

**Integration of Ngn1/Ngn2 cassette into iPSCs**

iPSCs were split and plated into a vitronectin-coated 6 well plate as described above, at a density such that cells were 50-70% confluent in clumps of 2-5 cells at the time of transfection. Following plating, cells were incubated overnight in E8 media with ROCK inhibitor (Fisher BDB562822), and changed into fresh E8 media the following morning. Thirty minutes prior to transfection (~24h after plating or when the density was 50-70%), cells were changed into mTESR-1 media (Cell Technologies 85850) and then transfected with 2.5 µg of donor DNA and 1.25 µg of each targeting construct (Supplemental Table 3) using Lipofectamine Stem (Invitrogen STEM00003) according to the manufacturer’s instructions. The following morning, cells were changed into fresh E8 media. Media was changed daily, and cells were screened for red fluorescence. When the partially positive colonies reached 100-500 cells, they were carefully scraped/aspirated using a P200 pipet tip and transferred to a new vitronectin-coated dish. This process was repeated, enriching the fluorescent cells until a 100% fluorescent colony was identified. This was then relocated to a new dish, and expanded for future use. The Ngn1/2 integration cassette and accompanying targeting constructs were a gift from M. Ward.

**iNeuron differentiation**

**Day 0**. Induced pluripotent stem cells were washed in PBS and incubated in prewarmed accutase (Sigma A6964) at 37°C for 8m. Four volumes of E8 media were added to the plate, and the cells were collected and pelleted at 200xg for 5m. The media was aspirated, and the pellet was resuspended in 1ml of fresh E8 media. Cells were counted using a hemocytometer, diluted, plated at a density of 20,000 cells/ml in E8 media with ROCK inhibitor and incubated at 37°C overnight. **Day 1.** Media was changed to N2 media (1x N2 Supplement (Gibco 17502-048), 1x NEAA Supplement (Gibco 11140-050), 10 ng/ml BDNF (Peprotech 450-02), 10 ng/ml NT3 (Peprotech 450-03), 0.2 µg/ml laminin (Sigma L2020), 2 mg/ml doxycycline (Sigma D3447) in E8 media). **Day 2.** Media was changed to transition media ((1x N2 Supplement, 1x NEAA Supplement, 10 ng/ml BDNF, 10 ng/ml NT3, 0.2 µg/ml laminin, 2 mg/ml doxycycline in half E8 media, half DMEM F12 (Gibco 11320-033)). **Day 3.** Media was changed into B27 media (1x B27 Supplement (Gibco 17504-044), 1x Glutamax Supplement (Gibco 35050-061), 10 ng/ml BDNF, 10 ng/ml NT3, 0.2 µg/ml laminin, 2 mg/ml doxycycline, and 1x Culture One (Gibco A33202-01) in Neurobasal-A (Gibco 12349-015)). **Day 6.** An equal volume of B27 media without Culture One was added to each well. **Day 9-21.** All cultures underwent a half-media change every 3d in fresh B27 media.

**Immunocytochemistry**

Neurons were fixed with 4% paraformaldehyde (PFA; Sigma P6148), rinsed with PBS, and permeabilized with 0.1% Triton X-100 (Bio-rad 161-0407) in PBS. Neurons were then treated with 10 mM glycine (Fisher BP381-1) in PBS, and incubated in a blocking solution (0.1% Triton X-100, 2% fetal calf serum (Sigma F4135), and 3% bovine serum albumin (BSA, Fisher BP9703-100) in PBS) at room temperature for 1h before incubation overnight at 4°C in primary antibody diluted in blocking buffer (Supplemental Table 4). Cells were then washed 3x in PBS and incubated at room temperature with Alexa Fluor 488 goat anti-rabbit (Life Technologies A11034), Alexa Fluor goat anti-mouse 594 (Life Technologies A11032), or Alexa Fluor donkey anti-rabbit 647 (Life Technologies A31573) secondary antibody diluted 1:500 in blocking solution for 1h. Following 3x washes in PBS containing 1:10000 Hoechst 33258 dye (Invitrogen H3569), neurons were imaged via fluorescence microscopy. High resolution images were obtained on a Zeiss LSM 800 with a

63x NA1.4 Oil/DIS Plan-Apochromat objective. Excitation was accomplished using 405, 488, 561, and 633 nm lasers.

**Modulation of neuronal activity**

Half of the existing media was removed from mature human iNeurons and replaced with fresh media and drug such that the final concentration on the cells was 4 mM tetraethylammonium chloride (TEA, Sigma T2265), 2 µM tetrodotoxin citrate (TTX, R&D Systems 1078), 2.5-5 µM of bicuculline (Sigma 14343), or 2.5-5 µM glutamate (Sigma G1251) alongside a volume-matched vehicle control. Cells were incubated at 37° C for 48h, then fixed, imaged, or harvested as needed.

**Monitoring calcium transients**

Mature iNeurons—differentiated as previously described from an iPSC line stably expressing gCaMP6f and mCherry (3)—were imaged for 100ms at 200ms intervals for a total of 100 frames, for a cumulative a 20s observation window. One location was imaged per well for 2-30 instances over a 6-12h period. Each neuron was identified as a region of interest using mCherry fluorescence, and the intensity of gCaMP6f signal was plotted over time. Individual traces were corrected for photobleaching, normalized to the median of each imaging period, and filtered for peaks below a discrete threshold to aid in spike identification. The number of peaks for each neuron and each imaging period was manually counted using a custom-designed graphical user interface. Events per second were averaged for each cell and compared across groups.

**CRISPR/Cas9 editing of iPSCs**

Oligos complementary to the target region (Supplemental Table 3) were annealed, digested, and ligated into the *BbsI* site in pX335-U6-Chimeric_BB-CBh-hSpCas9n(D10A) (Addgene #42335, deposited by Feng Zhang) or pX330S-4 (Addgene #58780, deposited by Feng Zhang) according to the protocol available from Addgene. iPSCs stably expressing Ngn1-2 under a dox-inducible promoter were split and transfected as described above with pX335 vectors encoding Cas9(D10A) and sgRNA pairs targeting sequences flanking the *TARDBP* start codon for D2-TDP43 or stop codon for TDP43-D2. Cells were cotransfected with the appropriate HDR vector encoding the Dendra2 open reading frame flanked by 400 bp of sequence homologous to that surrounding the *TARDBP* start codon (D2-TDP43) or stop codon (TDP43-D2) (in pUC-minus(M), synthesized by Blue Heron, LLC). Fluorescent cells were selected and successively passaged as described above to generate iPSC colonies in which 100% of cells expressed Dendra2-labeled TDP43.

**qRT-PCR**

RNA was isolated using the RNeasy Mini Kit (Qiagen 74106), and cDNA was reverse transcribed from 1 ug of the resultant RNA with the Bio-Rad iScript kit (Bio-Rad 170-8891) in a reaction volume of 20 µl. 0.5 µl of cDNA was used for each reaction as a template for quantitative (q)PCR, which was performed using Power SYBR Green (Applied Biosystems A25742) using the primers listed in Supplemental Table 5.

**Plasmids**

Plasmids pGW1-EGFP(1) (4), pGW1-TDP43-EGFP (5), and pGW1-mApple (5) were used both as experimental controls and to generate additional constructs (Supplemental Table 6).

To generate pGW1-sTDP43-EGFP, a geneblock comprised of the sTPD43-1 open reading frame (ORF) flanked by ApaI and AgeI restriction enzyme sites was generated by Integrated DNA Technologies (IDT). This geneblock was digested with ApaI and AgeI and cloned into the corresponding sites immediately upstream of the EGFP ORF in pGW1-EGFP(1).

To create pGW1-sTDP43(mNES)-EGFP, the sTDP43 open reading frame was amplified using a reverse primer to mutate the putative NES into five sequential glycine resides. The resulting amplicon was digested with ApaI and AgeI and cloned into corresponding sites in pGW1-EGFP(1).

To generate pGW1-EGFP(2), the EGFP open reading frame was PCR amplified from pGW1-EGFP(1). The resultant amplicon was digested with HindIII and Kpn1 restriction enzymes and cloned into the corresponding sites in pGW1-CMV.

To generate pGW1-EGFP-tail, sense and antisense oligomers with the sequence of the 18-amino acid tail were generated by IDT, designed such that annealing would result in cohesive ends identical to cut KpnI and NheI restriction enzyme sites. The annealed oligo was cloned into corresponding sites immediately downstream of the EGFP ORF in pGW1-EGFP(2).

To generate pGW1-EGFP-tail(mNES), sense and antisense oligomers with the sequence of the 18-amino acid tail in which the putative NES was replaced by 5 glycine residues were generated by IDT, designed such that annealing would result in cohesive ends identical to cut KpnI and NheI restriction enzyme sites. The annealed oligo was cloned into corresponding sites immediately downstream of the EGFP ORF in pGW1-EGFP(2).

To create pGW1-EGFP-TDP43, the TDP43 ORF was PCR amplified from pGW1-TDP43-EGFP. The resultant amplicon was digested with KpnI and NheI restriction enzymes and cloned into the corresponding sites immediately downstream of the EGFP ORF in pGW1-EGFP(2).

To generate pGW1-EGFP-sTDP43, the sTDP43-1 ORF was PCR amplified from pGW1-sTDP43-EGFP. The resultant amplicon was digested with KpnI and NheI restriction enzymes and cloned into the corresponding sites immediately downstream of the EGFP ORF in pGW1-EGFP(2).

To create pGW1-Halo, the HaloTag ORF was PCR amplified from pFN21A HaloTag PUM2 RBD R6SYE (a gift from A. Goldstrohm). The resultant amplicon was digested with XbaI and SbfI restriction enzymes and cloned into the corresponding sites in pGW1-CMV.

To create pGW1-TDP43-Halo, the TDP43 ORF was PCR amplified from pGW1-TDP43-EGFP. The resultant amplicon was digested with NheI and AgeI and cloned into corresponding sites in pGW1 to make pGW1-TDP43. The HaloTag ORF was then amplified from pFN21A HaloTag PUM2 RBD R6SYE, digested with XbaI and SbfI restriction enzyme sites and cloned into the corresponding sites immediately downstream of the TDP43 ORF in pGW1-TDP43.

To generate pGW1-sTDP43-Halo, the sTDP43-1 ORF was PCR amplified from pGW1-sTDP43-EGFP. The resultant amplicon was digested with AgeI and NheI and cloned into corresponding sites immediately upstream of the HaloTag ORF in pGW1-Halo.

To generate the TDP43 autoregulatory reporter(5), a 3 kb segment extending from *TARDBP* exon 6 to the 3' UTR was amplified from genomic DNA. The resultant amplicon was digested with BsrGI and SfiI and cloned into corresponding sites immediately downstream of the mCherry ORF in pCAGGs-mCherry.

Shuttle-RFP (pcDNA3.1-NLS-mCherry-NES) was purchased from Addgene (#72660, donated by B. Di Ventura and R. Eils). The CFTR minigene reporter was a gift from Y. Ayala, and pCaggs-mCherry and pGW1-CMV were gifts from S. Finkbeiner. The shTARDBP and non-targeting shRNA constructs were purchased from Dharmacon (V3SH11240-224779127, VSC11712).

All constructs were verified by Sanger sequencing and described in Supplemental Table 6.

**Primary neuron cell culture and transfection**

Cortices from embryonic day (E)19-20 Long-Evans rat embryos were dissected and disassociated, and primary neurons were plated at a density of $6x10^5$ cells/ml in 96-well plates, as described previously (6). At *in vitro* day (DIV) 4, neurons were transfected with 100 ng pGW1-mApple to mark cell bodies and 50-100 ng of an experimental construct using Lipofectamine 2000 (Invitrogen 52887), as previously described (5, 7, 8). Following transfection, cells were placed in either Neurobasal Complete Media (Neurobasal (Gibco 21103-049), 1x B27, 1x Glutamax, 100 units/mL Pen Strep (Gibco 15140-122)) or NEUMO photostable medium with SOS (Cell Guidance Systems M07-500) and incubated at 37°C in 5% $CO_2$.

**Longitudinal fluorescence microscopy and automated image analysis**

Neurons were imaged as described previously (9, 10) using a Nikon Eclipse Ti inverted microscope with PerfectFocus3a 20X objective lens and either an Andor iXon3 897 EMCCD camera or Andor Zyla4.2 (+) sCMOS camera. A Lambda XL Xenon lamp (Sutter) with 5 mm liquid light guide (Sutter) was used to illuminate samples, and custom scripts written in Beanshell for use in μManager controlled all stage movements, shutters, and filters. Custom ImageJ/Fiji macros and Python scripts were used to identify neurons and draw both cellular and nuclear regions of interest (ROIs) based upon size, morphology, and fluorescence intensity. Fluorescence intensity of labeled proteins was used to determine protein localization or abundance. Custom Python scripts were used to track ROIs over time, and cell death marked a set of criteria that include rounding of the soma, loss of fluorescence and degeneration of neuritic processes (8).

**Culturing and transfecting HEK293Ts**

Human embryonic kidney (HEK) 293T cells were cultured in DMEM (GIBCO 11995065), 10% FBS (Gibco ILT10082147), 1x Glutamax, and 100 units/mL Pen Strep at 37°C in 5% $CO_2$ (Supplemental Table 7). HEK293T cells are originally female in origin, are easily transfected, and have been transformed with SV40 T-antigen. HEK293T cells were transfected with Lipofectamine 2000 according to the manufacturer's instructions.

**Immunoprecipitation using HaloLink**

HEK293T cells were transfected with Halo-tagged constructs of interest. Two days after transfection, cells were collected in PBS and pelleted at 21,000xg for 5m. The cells were then resuspended in 100 µl lysis buffer (50 mM Tris-HCl, 150 mM NaCl, 1% Triton X-100, 0.1% sodium deoxycholate). After incubation on ice for 15m, cells were passed through a 27.5 G needle and pelleted at 21,000xg for 10m at 4°C. 100 µg of protein was then added to 100 µl of prewashed HaloLink resin (Promega G1914), which was prepared by washing and pelleting for 2m at 800xg 3x in wash buffer (100 mM Tris pH 7.5, 150 mM NaCl, 1 mg/ml BSA, 0.005% IGPAL). Sufficient wash buffer was added to ensure an equal volume for all conditions (~400 µl), and samples were incubated on a tube rotator for 30m at room temperature. Samples were then pelleted at 800xg for 2m, saving the supernatant. The beads were then washed 3x in wash buffer, and resuspended in elution buffer (1% SDS, 50 mM Tris-HCl pH 7.5) and 10x sample buffer (10% SDS, 20% glycerol, 0.0025% bromophenol blue, 100 mM EDTA, 1 M DTT, 20 mM Tris, pH 8). Samples were then boiled at 95° C for 10m, and loaded onto a 10% SDS-PAGE gel alongside 10 µg of input protein and a fixed volume of supernatant to assess binding efficiency. The gel was run at 120 V, and samples were then transferred at 100 V at 4°C onto an activated 2 um polyvinylidene difluoride (PVDF) membrane (Bio-Rad 1620177), blocked with 3% BSA in 0.2% Tween-20 (Sigma P9614) in Tris-buffered saline (TBST) for 1h, and blotted overnight at 4°C with primary antibody in 3% BSA in TBST (Supplemental Table 4). The following day, blots were washed in 3x in TBST, incubated at room temperature for 1h with donkey anti-mouse 680 RD (Li-Cor 926-68072) and donkey anti-rabbit 800 CW (Li-Cor 925-32213) secondary antibodies, both diluted 1:5,000 in 3% BSA in TBST. Following treatment with secondary antibody, blots were washed 3x in TBST, placed in Tris-buffered saline, and imaged using an Odyssey CLx Imaging System (LI-COR).

**Differential solubility fractionation**

HEK293T cells were transfected in a 6-well plate with 3 µg of DNA/well using Lipofectamine 2000 according to the manufacturer’s instructions. Two days after transfection, cells were collected in PBS and pelleted at 21,000xg for 5m. Cells were then resuspended in RIPA buffer (Thermo Scientific 89900) with protease inhibitors (Roche 11836170001) and incubated on ice for 15m. Lysates were then sonicated at 80% amplitude with 5s on/5s off for a total of 2m using a Fisher Brand Model 505 Sonic Dismembrenator (ThermoFisher). Samples were centrifuged at 21,000xg for 15m at 4°C, after which the supernatant was removed and saved as the RIPA-soluble fraction. The RIPA-insoluble pellet was washed in RIPA once more and resuspended in urea buffer (7 M urea, 2 M thiourea, 4% CHAPS, 30 mM Tris, pH 8.5) and incubated on ice for 5m. Samples were then centrifuged at 21,000xg for 15m at 4°C, and the supernatant was saved as the RIPA-insoluble, urea-soluble fraction. The RIPA-soluble samples were quantified and 10-30 µg of protein/well was diluted in RIPA buffer with 10x sample buffer. For urea fractions, equal volumes of each sample across conditions was diluted in urea buffer and 10x sample buffer. The RIPA-soluble samples were boiled for 10m before 10-30 µg of all samples were loaded onto a 10% SDS-PAGE gel with stacking gel and run at 120 V. The blot was then transferred and probed as described above.

**RNA sequencing**

Raw reads from murine frontal cortex and spinal cord (11–13), as well as human spinal cord, frontal cortex and cerebellum (11–15), were downloaded from Gene Expression Omnibus (GEO) with the SRA Toolkit v2.9.2. Reads were trimmed with TrimGalore v0.6.0 using automatic adapter detection and a minimum Phred score of 20. For alignment-free transcript-level quantification, trimmed reads were quantified using Salmon v0.13.1 (Patro, 2017) and imported into the RStudio using txImport v1.12.0 (Soneson, 2015) to generate transcript-level summaries (16,17). The Ensembl genome assemblies and transcript annotations from GRCh38.96 and GRcm38.96 were used as human and mouse references, respectively. For alignment-based analysis of mouse datasets, trimmed reads were aligned with hisat2 v2.0.5 and raw counts were quantified for unique splice donor/acceptor combinations present in unique *TARDBP* isoforms. In each case, splicing events were visualized using IG Viewer software (Broad Institute).

**TDP43 knockdown in N2A cells**

N2A mouse neuroblastoma cells were cultured in DMEM (GIBCO 11995065), 10% FBS (Gibco ILT10082147), 1x Glutamax, and 100 units/mL penicillin/streptomycin at 37°C in 5% $CO_2$ (Supplemental Table 7). Cells were transfected using Lipofectamine 2000 (ThermoFisher) according to the manufacturer's instructions, with artificial microRNAs (amiRNAs) directed against *TARDBP* or a scrambled control (18). Transfected N2As were incubated for 72h, harvested, and immunoblotted to verify TDP43 knockdown.

**Tissue preparation and immunohistochemistry in murine tissue**

Vertebral columns were dissected from 5 month old C57Bl6 J mice, fixed in 4% paraformaldehyde (PFA) at 4°C for 48h, washed in phosphate buffered saline (PBS), and dissected to extract the spinal cords. The lumbar enlargement was sub-dissected, cryoprotected in 30% sucrose at 4°C, embedded in M1 matrix (Thermo Scientific #1310) in a silicone mold, frozen on dry ice, and sectioned at a thickness of 16µm onto charged slides (Thermo Scientific J1800AMNZ). Sections were then briefly air dried and stored at -80°C. For immunohistochemistry (IHC), sections were washed in distilled water, and blocked and permeabilized in blocking buffer (5% BSA, 0.5% Triton X-100, and 5% goat serum (Gibco 16210-064)) for 1h at room temperature. Slides were then incubated with primary antibody at 4°C overnight in blocking buffer diluted 2-fold with PBS (Supplemental Table 4). Sections were washed 3x for 5m in PBS, then incubated at room temperature for 1h with Alexa Fluor goat anti-mouse 488 (Life Technologies AB150113), Alexa Fluor donkey anti-goat 647 (Life Technologies AB150131), and Alexa Fluor goat anti-rabbit 568 (Life Technologies AB175470) secondary antibodies diluted 1:500 in blocking solution. Sections were then washed 3x for 5m, counterstained, and mounted with Vectashield Hardset with DAPI (Vector Labs H-1500). Images were acquired using Olympus Whole Slide Scanner (VS120) with a 40x objective.

**Immunohistochemistry in human tissue**

Paraffin-embedded human cortex and spinal cord obtained from the University of Michigan Brain Bank were cut into 5 μm thick sections and mounted on glass slides. Tissue samples were photobleached prior to immunofluorescence (19). Briefly, slides were placed on ice, under a 7 Band Spectrum LED Light (HTG Supply 432W HTG-432-3W-7X) at 4°C for 12h. Slides were deparaffinized at 65°C for 20m, and rehydrated 5m sequentially in xylene (Fisher X3S-4), 100% ethanol (Fisher 3.8L), 95% ethanol, 70% ethanol, 50% ethanol, and PBS. Slides were then permeabilized with 0.1% Triton X-100 in PBS, and treated with 10 mM glycine in PBS. They were then incubated in a blocking solution (0.1% Triton X-100, 2% fetal calf serum, and 3% BSA in PBS) at room temperature for 1h before incubation overnight at 4°C in primary antibody diluted in blocking buffer (Supplemental Table 4). Slides were then washed 3x in PBS and incubated at room temperature with Alexa Fluor 488 goat anti-rabbit (Life Technologies A11034), Alexa Fluor goat anti-mouse 594 (Life Technologies A11032), and/or Alexa Fluor goat anti-chicken 647 (Life Technologies A21449) secondary antibody diluted 1:500 in blocking solution for 1h. Following 3x washes in PBS containing 1:10,000 Hoechst 33258 dye (Invitrogen H3569), slides were mounted in mounting media (Fisher SP15-500) and allowed to dry in the dark overnight before being imaged the following day. Images were acquired using a Nikon Microphot-FXA microscope (Nikon, 1985) in combination with a 60x oil-immersion objective, a QIClick CCD Camera (Q Imaging, 7400-82-A1), and an X-Cite Series 120 light source (Lumen Dynamics).

**Differentiation of iPSC-derived astrocytes**

iPSC-derived astrocyte progenitors were derived, cultured, and expanded as described previously (20) (Supplemental Table 7). The glial progenitors were pre-differentiated in Neurobasal Medium containing 1% B27 Supplement, 1% NEAA, and 1% PS for 10d on 6-well plates before cryopreservation. Before the experiment, batches of pre-differentiated astrocytes were defrosted and plated at 50k per well in 8-well Ibidi imaging chambers (Ibidi 80841), and cultured for an additional 6 days in Neurobasal Medium containing 1% B27, 1% NEAA, 1% PS, and 20 ng/mL ciliary neurotrophic factor (CNTF, Thermo Fisher PHC7015). Astrocytes were then fixed in 4% PFA for 15m and immunostained as described above (Supplemental Table 4) using Alexa Fluor goat anti-rabbit 647 (Thermo Fisher A-21245) and Alexa Fluor goat anti-mouse IgG1 (Thermo Fisher A-21121) secondary antibodies diluted 1:1000 in 3% goat serum in PBS. Imaging was performed on a Leica DMi8 with CooLED light source at 63x and analyzed with ImageJ.

**Statistical analysis**

Statistical analyses were performed in R or Graphpad Prism 7. For primary neuron survival analysis, the open-source R survival package was used to determine hazard ratios describing the relative survival between conditions through Cox proportional hazards analysis (8). Significance determined via the two-tailed t-test was used to assess differences between treatment groups for neuronal activity and transcript abundance via RT-PCR. The Kolmogorov-Smirnov test was used to assess differences between the distribution of TDP43 abundance in neurons under different activity conditions. One-way ANOVA with Tukey's or Dunnett's post-tests were used to assess significant differences among nuclear/cytoplasmic ratios, binding affinity, TDP43 splicing activity, and TDP43 autoregulation. Data are shown as mean ± SEM unless otherwise stated.

**Supplemental Figures**

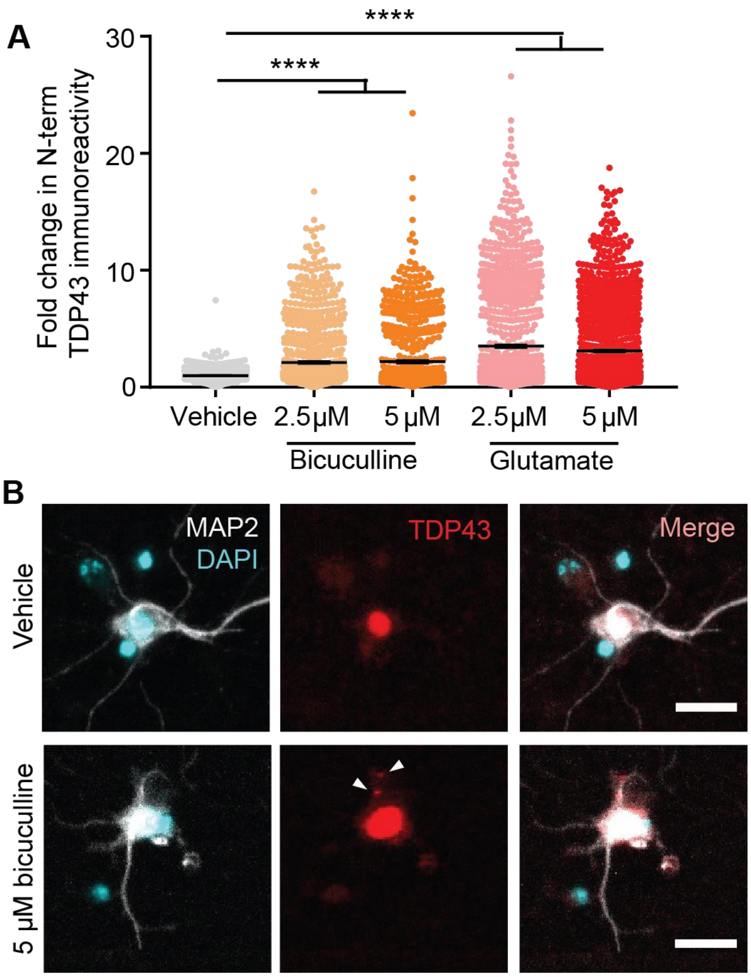


**Supplemental Figure 1. Multiple drivers of neuronal hyperexcitability upregulate N-terminal TDP43.**

(**A**) DIV 28 rodent primary mixed cortical neurons treated with bicuculline (orange) or glutamate (red) show an increase in N-terminal TDP43 immunoreactivity compared to a vehicle-treated control (gray) via ICC. Vehicle n=879, 2.5 μM bicuculline n=1166, 5 μM bicuculline n=837, 2.5 μM glutamate n=1315, 5 μM bicuculline n=1536, data represent two replicates, ****$p<0.0001$, one-way ANOVA with Dunnett's post-test. (**B**) Representative images of TDP43 staining in vehicle- or bicuculline-treated neurons. White arrows indicate cytosolic puncta. Scale bar in (**B**), 20 μM.

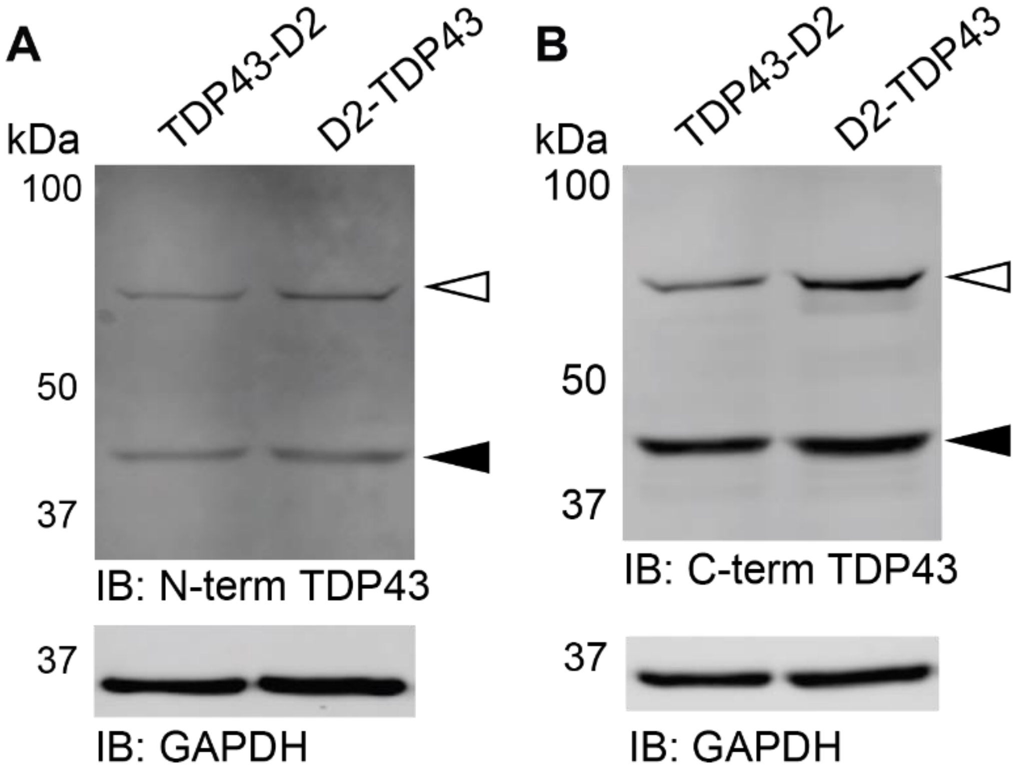


**Supplemental Figure 2. Validation of Dendra2-tagged iPSC lines.**
iPSCs immunoblotted for both (**A**) N-terminal and (**B**) C-terminal TDP43 indicate that both iPSCs lines are heterozygous for the insertion of Dendra2 (white arrow). Black arrow indicates untagged TDP43, GAPDH served as a loading control.

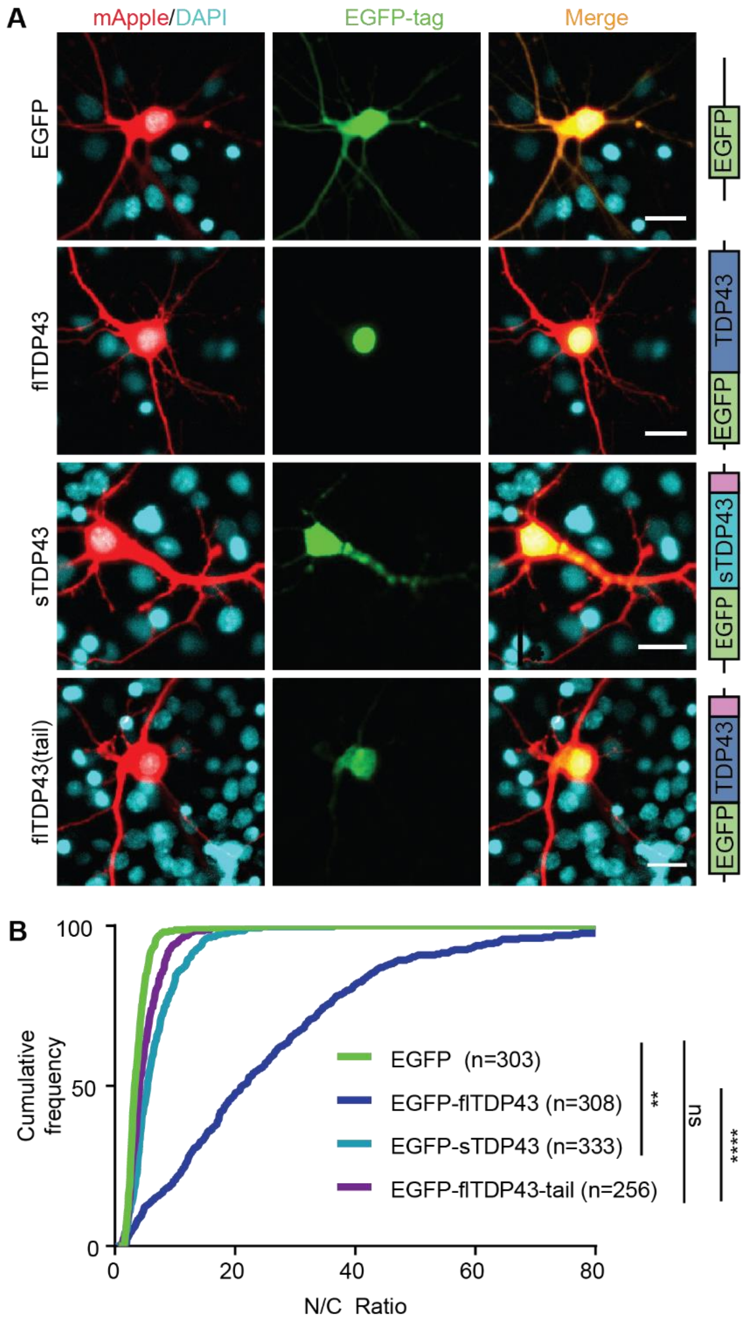


**Supplemental Figure 3. The sTDP43 C-terminal tail shifts flTDP43 localization to the cytoplasm.**

(**A**) Rodent primary mixed cortical neurons were transfected with mApple and EGFP-tagged TDP43 isoforms, then imaged by fluorescence microscopy. (**B**) Much like EGFP-sTDP43, EGFP-flTDP43-tail is significantly more cytosolic than EGFP-flTDP43. EGFP n=303, EGFP-flTDP43 n=308, EGFP-sTDP43 n=333, EGFP-flTDP43-tail n=256, consistent among 3 replicates, **p<0.01, ****p<0.0001, one-way ANOVA with Dunnett's post-test. Scale bar in (**A**), 20 µm.

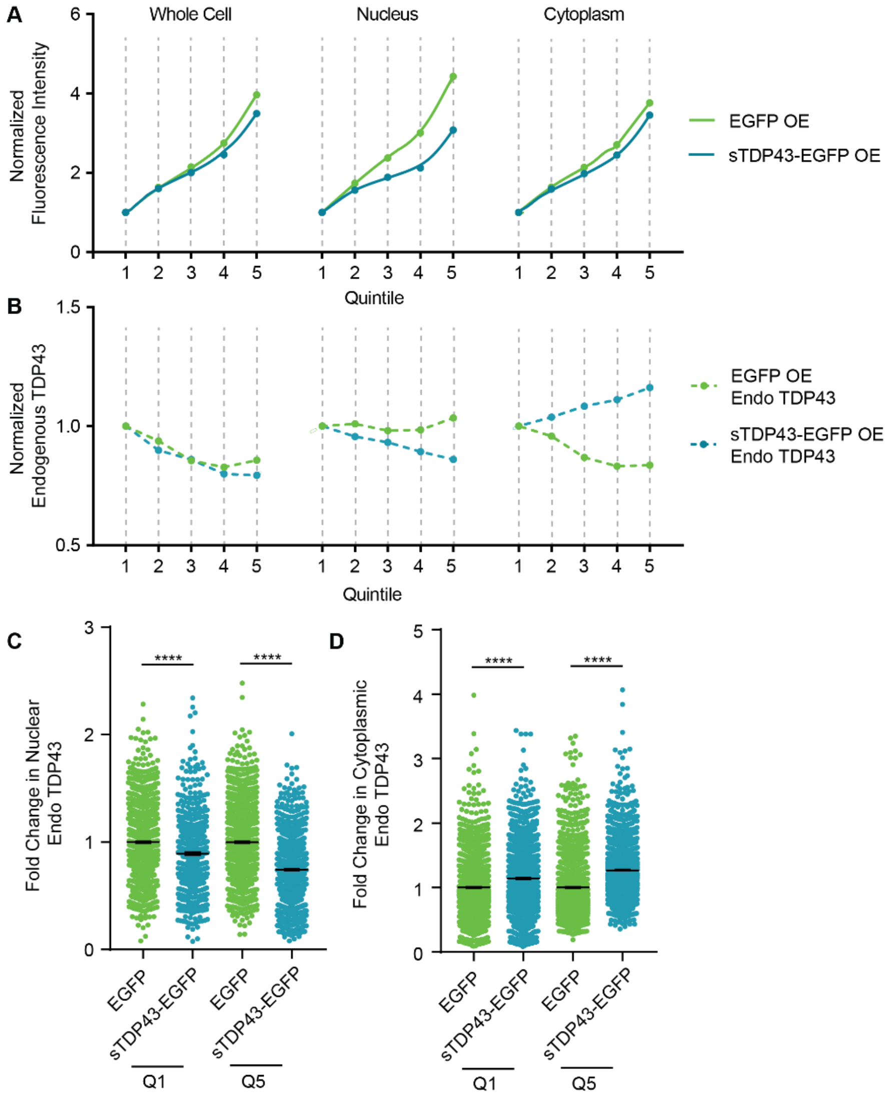


**Supplemental Figure 4. sTDP43 drives endogenous TDP43 mislocalization in a dose-dependent manner.** (**A**) HEK293T cells were transfected with EGFP or EGFP-tagged sTDP43, then immunostained using an antibody that recognizes the C-terminus of endogenous TDP43. Whole cell ROIs were determined by mApple fluorescence, and nuclear ROIs were determined by DAPI staining. Cells were sorted into quintiles based on EGFP fluorescence. (**B**) Endo TDP43 signal is comparable between cells overexpressing EGFP and sTDP43-EGFP at the whole cell level. However, nuclear endo TDP43 is reduced in a dose-dependent manner in cells overexpressing sTDP43-EGFP while cytosolic endo TDP43 increases in a dose-dependent manner. (**C**) Both low (Q1) and high (Q5) expression of sTDP43-EGFP results in a reduction in nuclear endo TDP43. Q1 EGFP n=1117, Q1 sTDP43-EGFP n=638, Q5 EGFP n=1203, Q5 n=1005, consistent between two replicates, ****$p<0.0001$, two-tailed t-test. (**D**) Cytoplasmic endogenous TDP43 is elevated by the expression of sTDP43-EGFP in both low (Q1) and high (Q5) expressing cells. Q1 EGFP n=1599, Q1 sTDP43-EGFP n=1599, Q5 EGFP n=1419, Q5 sTDP43-EGFP n=1492, consistent between two replicates, ****$p<0.0001$, two-tailed t-test.

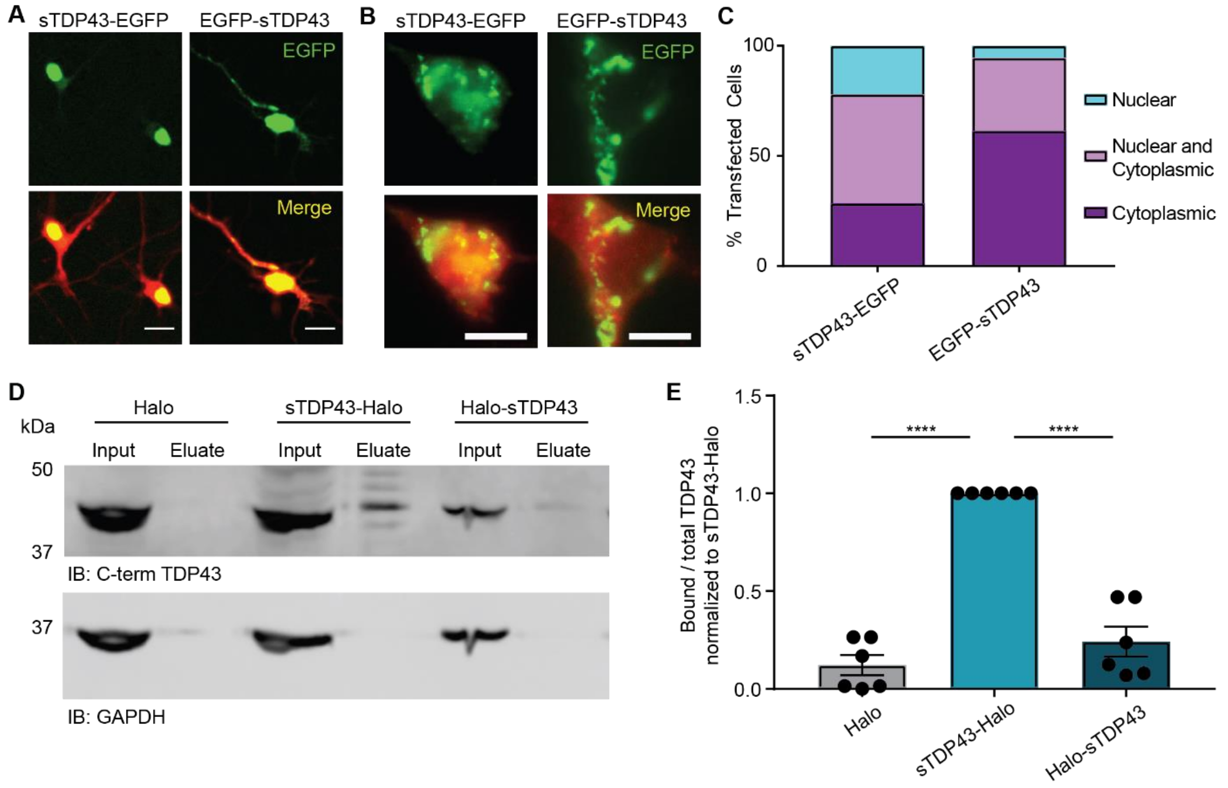


**Supplemental Figure 5. The position of exogenous protein tags influences sTDP43 localization and binding.**

(**A**) Fluorescence microscopy of rodent primary mixed cortical neurons or (**B**) HEK293T cells expressing N- or C-terminally tagged sTDP43 (green) as well as the whole cell marker mApple (red). (**C**) Characterization of sTDP43 localization in HEK293T cells expressing C- or N-terminally tagged sTDP43. sTDP43-EGFP n=196, EGFP-sTDP43 n=200. (**D**) sTDP43 fused with an N- or C-terminal HaloTag was expressed in HEK293T cells and immunoprecipitated with HaloLink resin. Bound, endogenous TDP43 was immunoblotted with a C-terminal TDP43 antibody. GAPDH served as a loading control. Input, (I); eluate, (E). (**E**) Quantification of data shown in (**D**), demonstrating the fraction of total TDP43 bound to HaloTag-sTDP43, sTDP43-HaloTag, or HaloTag alone. Data were combined from 6 replicates, ****$p<0.0001$, one-way ANOVA with Dunnett's post-test. Scale bars in (**A**) 20 µm, (**B**) 10 µm.

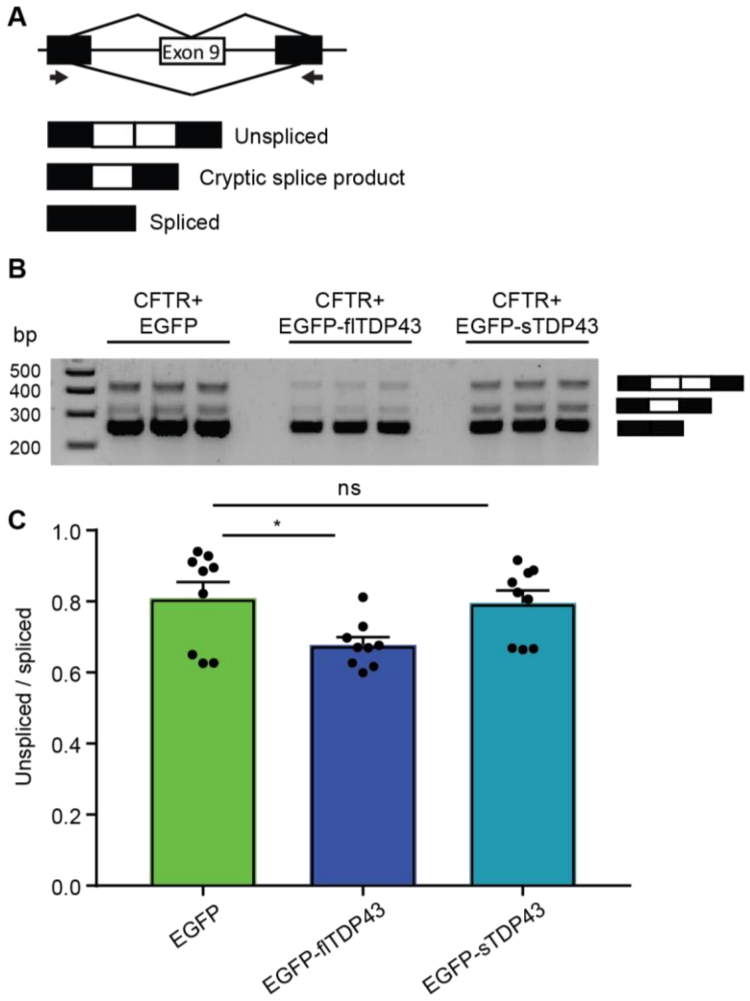


**Supplemental Figure 6. sTDP43 is deficient in splicing activity.**
(**A**) Schematic of the *CFTR* minigene reporter. TDP43-mediated splicing of the reporter results in exon 9 exclusion. Arrows indicate primers used to amplify the splice junction. (**B-C**) EGFP-flTDP43, but not EGFP-sTDP43, effectively excludes *CFTR* exon 9 in HEK293T cells overexpressing the reporter. 3 replicates, *p<0.05, one-way ANOVA with Dunnett's post-test.

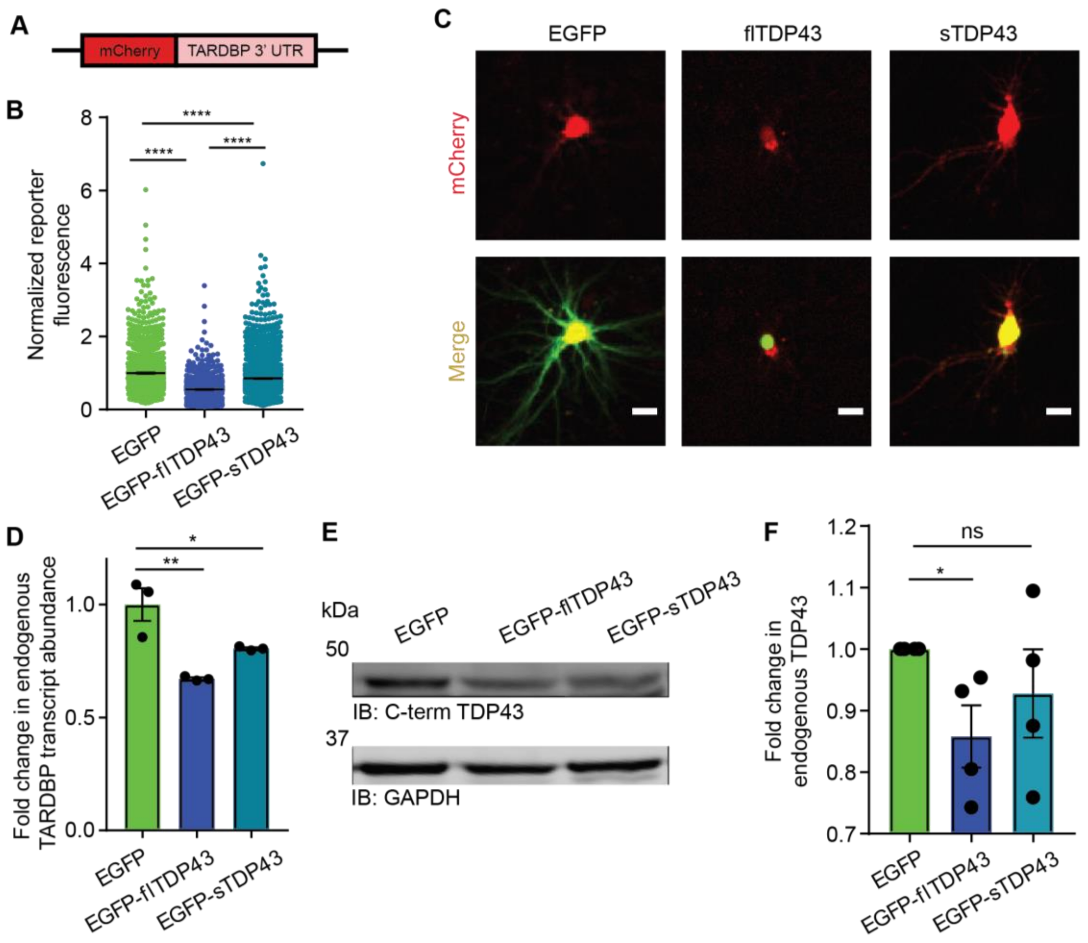


**Supplemental Figure 7. The autoregulatory function of sTDP43 is impaired.**
(**A**) Schematic of the TDP43 autoregulation reporter in which the fluorescent protein mCherry is fused to the *TARDBP* 3' UTR. (**B**) Rodent primary mixed cortical neurons overexpressing EGFP-flTDP43 show a significant reduction in reporter signal, consistent with autoregulation, while those overexpressing EGFP-sTDP43 do so to a lesser degree. EGFP n=2044, EGFP-flTDP43 n=2375, EGFP-sTDP43 n=2208, 3 replicates, ****$p<0.0001$, one-way ANOVA with Dunnett's post-test. (**C**) Fluorescence microscopy of rodent primary mixed cortical neurons demonstrating reduced reporter fluorescence in neurons co-expressing EGFP-flTDP43, in comparison to those co-expressing EGFP or EGFP-sTDP43. (**D**) qRT-PCR of HEK293T cells overexpressing EGFP, EGFP-flTDP43, or EGFP-sTDP43. Endogenous full-length *TARDBP* transcript was detected using primers that flank the stop codon, and transcript levels were normalized to GAPDH. 3 replicates, *$p<0.05$, **$p<0.01$, one-way ANOVA with Dunnett's post-test. (**E**) HEK293T cells overexpressing each construct were immunoblotted for C-terminal TDP43, GAPDH served as a loading control. (**F**) Quantification of results shown in (E). 4 replicates, * $p<0.05$, one-way ANOVA with Dunnet's post-test. Scale bar in (**C**), 10 µm.

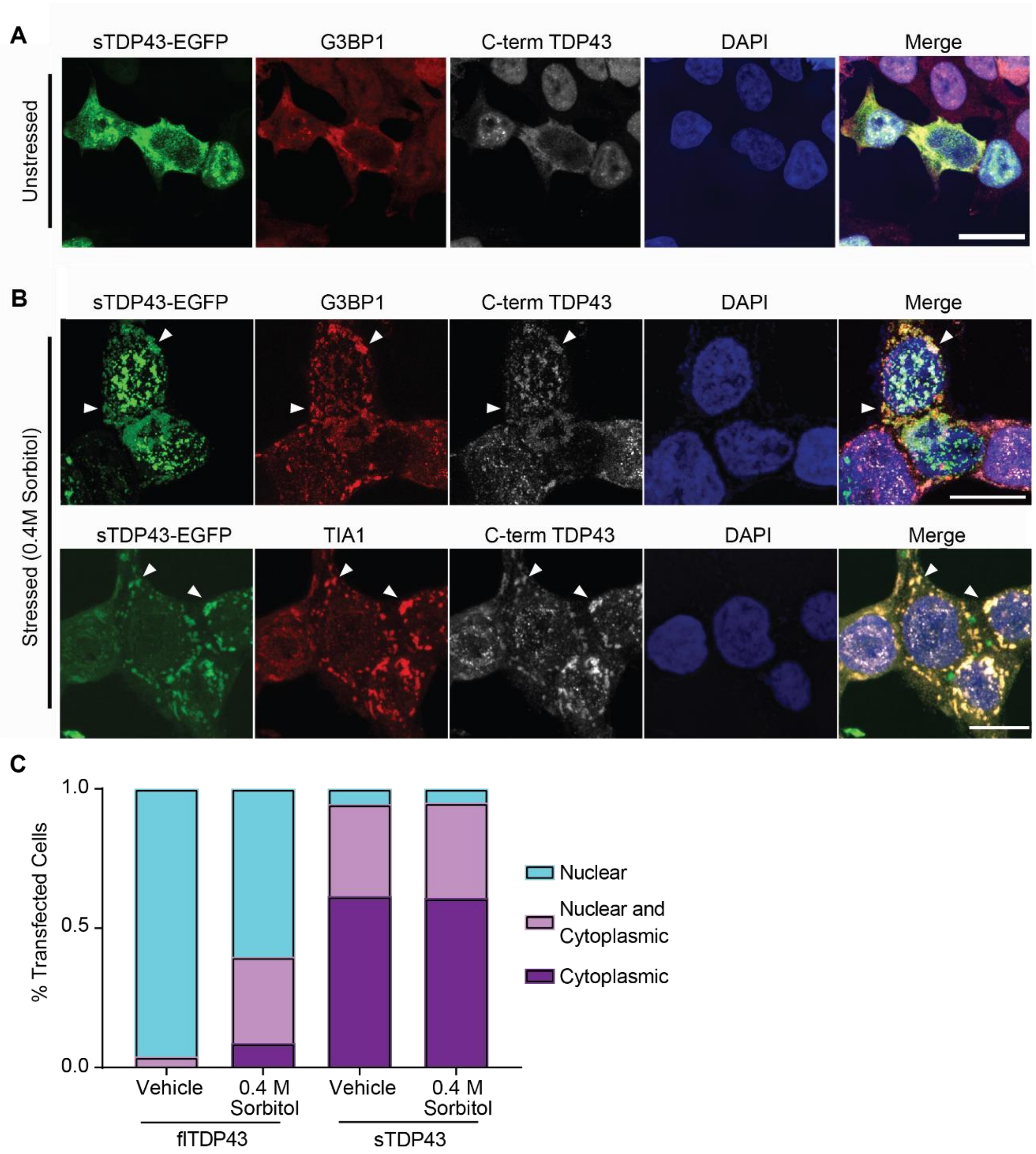


**Supplemental Figure 8. sTDP43 colocalizes with components of stress granules.**
(**A**) HEK293T cells were transfected with EGFP-tagged sTDP43 and immunostained using antibodies against the stress granule marker G3BP1 and endogenous TDP43. (**B**) Overexpressed sTDP43-EGFP colocalizes with endogenous TDP43 and stress granule markers G3BP1 and TIA1 in HEK293T cells treated with 0.4M sorbitol (arrows). (**C**) Assessment of stress-dependent changes in flTDP43 and sTDP43 localization. HEK293T cells were transfected with each construct and then stressed with sorbitol as before. The distribution of each protein was characterized as primarily nuclear, primarily cytoplasmic, or both for each cell. flTDP43 + vehicle n=216, flTDP43 + sorbitol n=219, sTDP43 + vehicle n=200, sTDP43 + sorbitol n=158. Scale bar in (**A**) 20 µm, scale bar in (**B**), 10 µm.

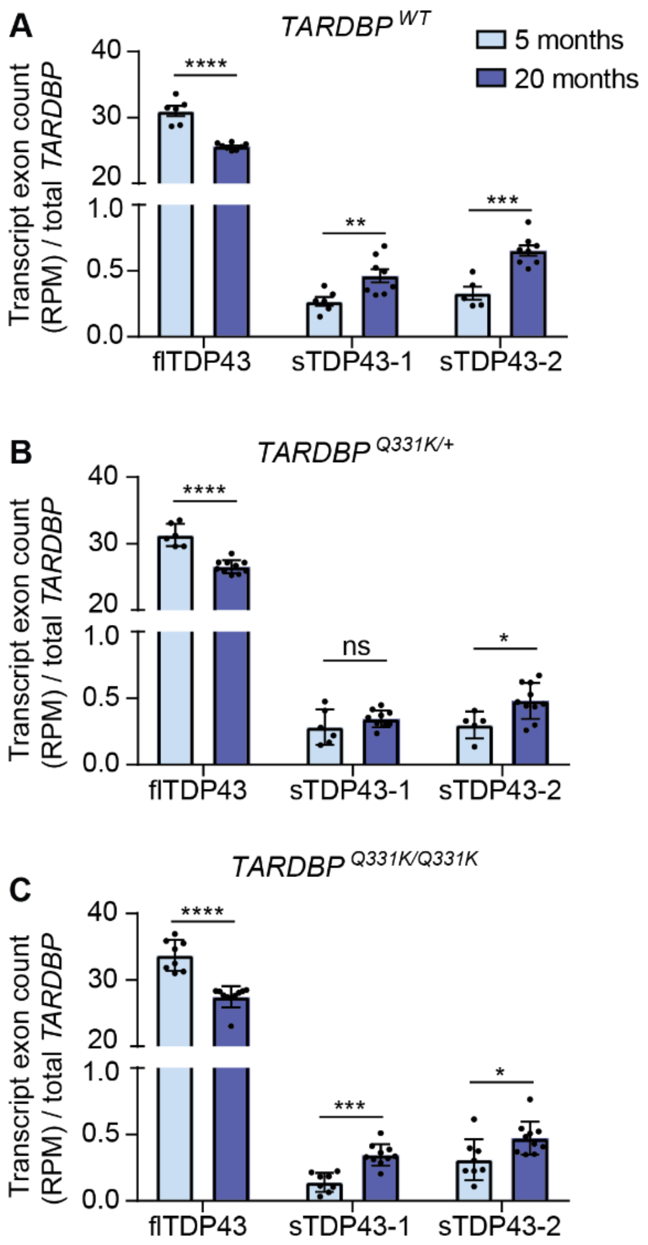


**Supplemental Figure 9. sTDP43 transcript abundance increases with age but is unaffected by the TDP43(Q331K) mutation.**

Murine frontal cortex collected at 5 (pale blue) or 20 (purple) months of age show both a decrease in flTDP43 and an increase in sTDP43-1 and -2 transcript abundance with age. This is observed in WT mice (**A**) as well as those that are hetero- (**B**) and homozygous (**C**) for *TARDBP*(Q331K) mutations. Graph depicts read counts normalized to reads per million for each library as a fraction of total *TARDBP*. 4 replicates, *$p<0.05$, **$p<0.01$, ***$p<0.001$, ****$p<0.0001$, multiple t-test with the Holm-Sidak correction.

**A**

| Original Study | Species | Library preparation | Read Length | Cell or Tissue Type | Disease Condition | n |
|---|---|---|---|---|---|---|
| White et al. 2018 | Mouse | Clontech SMART-seq and Illumina Nextera XT | 100 bp, paired | Laser captured spinal motor neuron | Wildtype | 4 |
| White et al. 2018 | Mouse | Illumina TruSeq | 100 bp, paired | Frontal cortex | Wildtype | 6 |
| Krach et al. 2018 | Human | NuGEN Ovation | 50 bp, single | Laser captured spinal motor neuron | Control and sALS | 21 |
| D'Erchia et al. 2017 | Human | Illumina TruSeq | 100 bp, paired | Ventral horn spinal cord | Control and sALS | 11 |
| Preduncio et al. 2015 | Human | Illumina TruSeq | 100 bp, paired | Cerebellum and Frontal cortex | Control, C9ALS, and sALS | 53 |

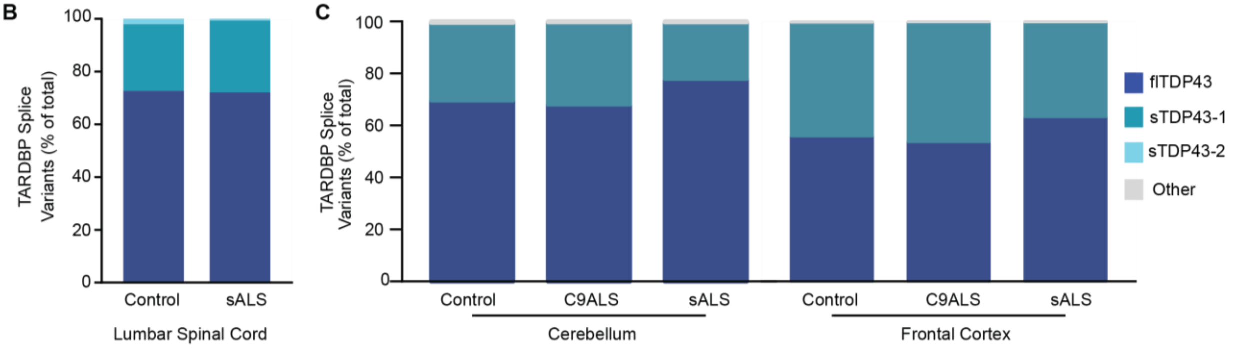


**Supplemental Figure 10. sTDP43 transcripts are present in a variety of tissue types and disease states.**

(**A**) Summary of previously published RNA-seq studies analyzed for sTDP43. (**B**) sTDP43-1 represents 30% of *TARDBP* transcripts in spinal cord ventral horn homogenate isolated from both control and sALS patients, with the remainder corresponding to flTDP43. (**C**) Similarly, sTDP43-1 makes up 30% of *TARDBP* transcripts in cerebellum, and 55% of *TARDBP* transcripts in frontal cortex. In each case, sTDP43-2 transcripts were largely undetectable, and there was no significant change in transcript isoform abundance in C9ALS or sALS patients compared to controls.

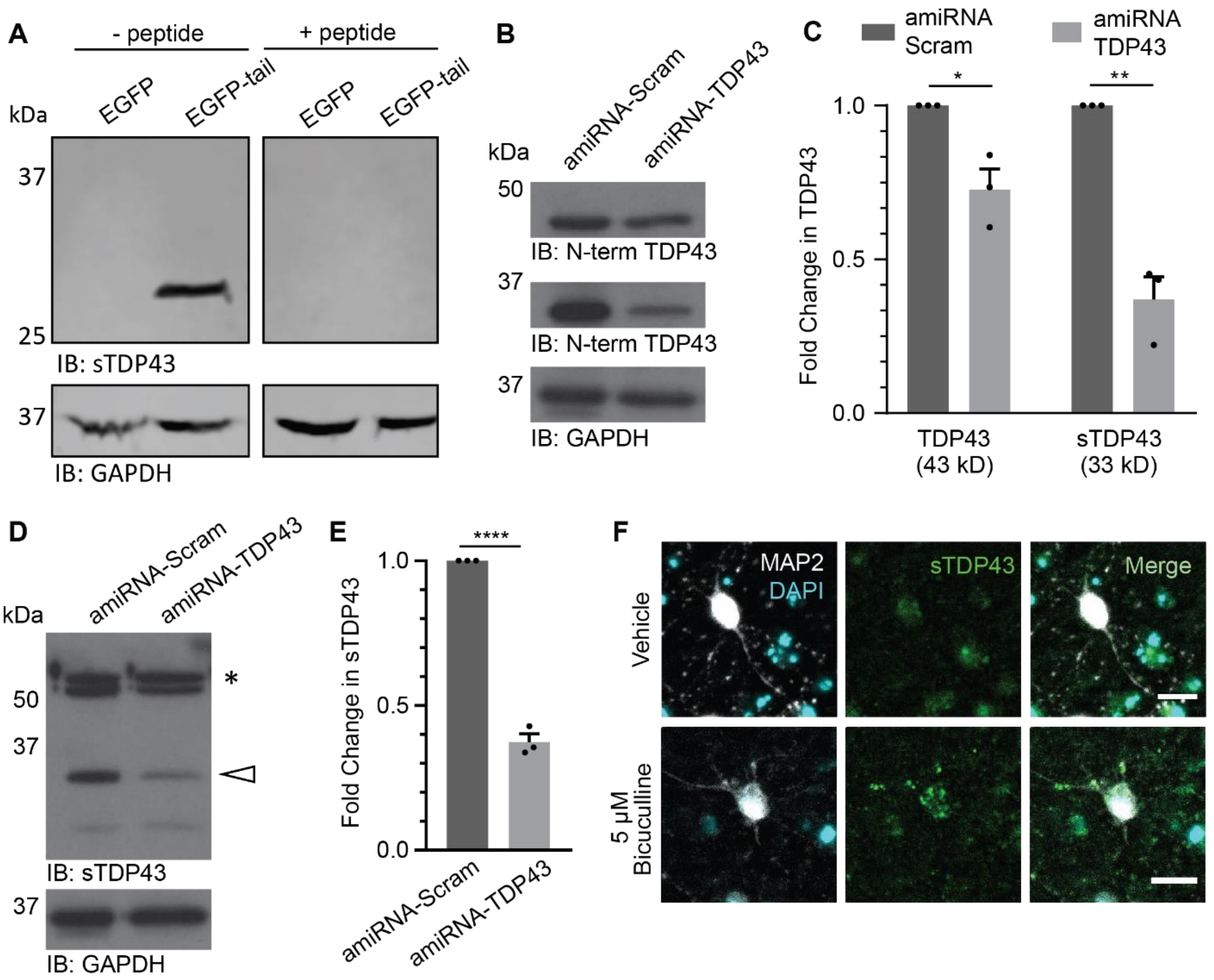


**Supplemental Figure 11. Validation of the sTDP43-specific antibody.**
(**A**) The sTDP43 antibody specifically recognizes EGFP fused to the 18-amino acid C-terminus of sTDP43, but not EGFP alone. Preincubation with a peptide corresponding to the sTDP43 C-terminal tail abolishes the signal. (**B**) N2A cells were transiently transfected with artificial microRNAs (amiRNAs) targeting TDP43, and proteins were separated by SDS-PAGE and immunoblotted using antibodies against N-terminal TDP43. Two bands were detected, the first at 43 kD corresponding to flTDP43 and the second at 33 kD corresponding to sTDP43. GAPDH served as a loading control. (**C**) Compared to scrambled amiRNAs, cells expressing amiRNA-TDP43 show a ~30% reduction of the 43kD species and a ~65% reduction of the 33kd species (3 replicates, *$p<0.05$, **$p<0.01$, two-tailed t-test). (**D**) Immunoblotting with the sTDP43-specific antibody detects a 33 kD band (white arrow), as well as a non-specific band at ~55 kD (asterisk). (**E**) sTDP43 shows a ~60% reduction in cells expressing amiRNA-TDP43 compared to scrambled control (3 replicates, ****$p<0.0001$, two-tailed t-test). (**F**) DIV 28 rodent primary mixed cortical neurons were treated with 5 μM bicuculline for 48h and immunostained for sTDP43. Bicuculline-treated neurons show cytosolic sTDP43 inclusions that are absent in vehicle-treated controls. Scale bar in (**F**), 20 μM.

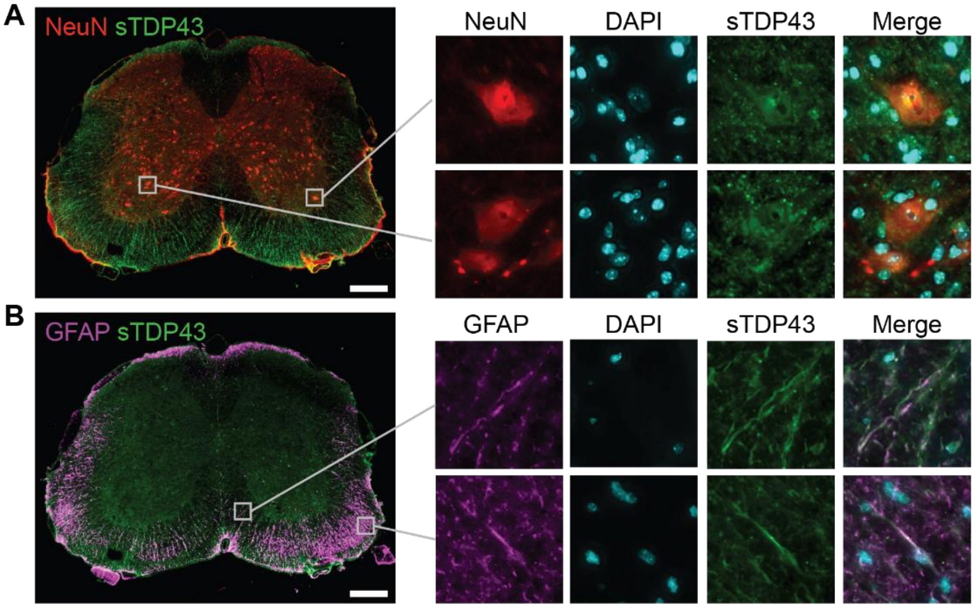


**Supplemental Figure 12. Endogenous sTDP43 is expressed by neurons and glia in murine lumbar spinal cord.**
Immunohistochemistry in murine spinal cord showing colocalization of sTDP43 immunoreactivity (green) with the neuronal marker NeuN (red) in the ventral horn (**A**) and with the astrocytic marker GFAP (**B**, purple). Scale bars (**A**) and (**B**), 200 µm.

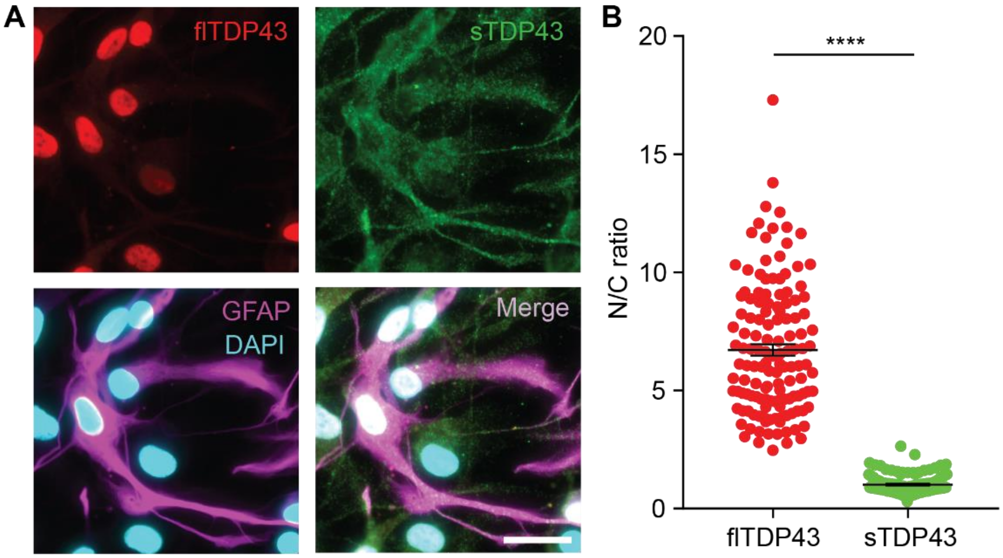


**Supplemental Figure 13. Endogenous sTDP43 is produced by human iPSC-derived astrocytes.**

(**A**) Immunocytochemistry using antibodies against flTDP43 (red) and sTDP43 (green) in astrocytes differentiated from human iPSCs. (**B**) Reflecting their unique subcellular distributions, flTDP43 displays a significantly higher nucleocytoplasmic (N/C) ratio than sTDP43 (3 replicates, flTDP43 n=136, sTDP43 n=136, ****p<0.0001, two-tailed t-test). Scale bar in (**A**) 10 µm.

**A**

| Case | Sex | Age | Age of Onset | PMI (Hours) | Clinical Diagnosis | Neuropath Diagnosis |
|---|---|---|---|---|---|---|
| CT-1 | Male | 56 | 49 | 14 | MSA-P | MSA |
| CT-2 | Male | 88 | NA | 5 | Normal | DLB with AD |
| ALS-1 | Female | 55 | 53 | Unknown | Bulbar Onset ALS | C9 ALS |
| ALS-2 | Female | 60 | Unknown | 33 | Limb Onset ALS | sALS |
| ALS-3 | Female | 87 | 77 | 5 | Bulbar Onset ALS | sALS |

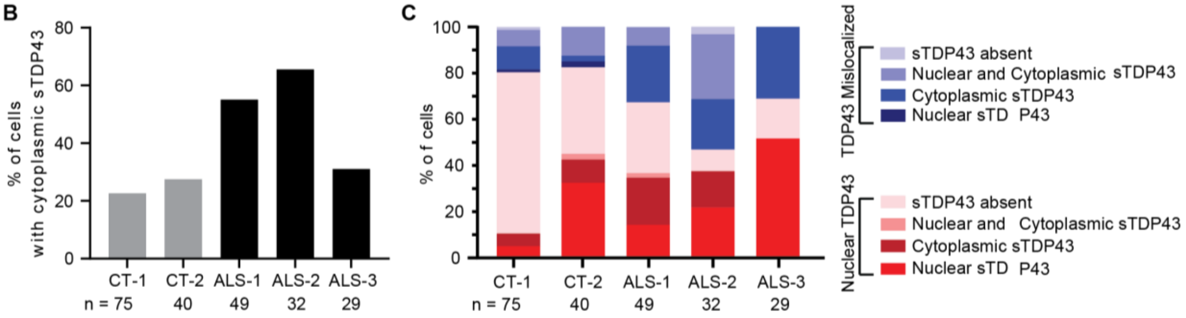


**Supplemental Figure 14. Characterization of sTDP43 pathology in ALS-patient tissue.** (**A**) Additional information for the control and ALS patient tissue used in these studies. PMI, post-mortem interval; MSA-P, multiple system atrophy with Parkinsonism; DLB with AD, Dementia with Lewy bodies with concurrent Alzheimer's Disease. (**B**) Percentage of cells with cytoplasmic sTDP43 identified in each control (gray) or ALS (black) patient. Number of cells counted per sample is listed below each column. (**C**) Characterization of sTDP43 localization for each patient. These data are further divided based on whether flTDP43 was nuclear (red) or cytoplasmically mislocalized (blue).

| Species | Isoform 1 Identity | Isoform 2 Identity |
| --- | --- | --- |
| Baboon (Papio anubis) | 99% | 99% |
| Drill (Mandrillus leucophaeus) | 96% | 96% |
| Angolan black and white colobus (Colobus angolensis palliatus) | 96% | 96% |
| Goats (Capra hircus) | 96% | 96% |
| Chinese tree shrew (Tupaia chinensis) | 96% | 96% |
| Horse (Equus caballus) | 96% | 96% |
| Cat (Felus catus) | 96% | 96% |
| Ground Squirrel (Ictidomys tridecemlineatus) | 96% | 96% |
| Shrew (Mus pahari) | 96% | 96% |
| Brown rat (Rattus novegicus ) | 96% | 96% |
| Deer mouse (Peromyscus maniculatus bairdii) | 96% | 96% |
| Prairie vole (Microtus ochrogaster) | 96% | 96% |
| House mouse (Mus musculus) | 96% | 96% |
| Ryukyu mouse (Mus caroli) | 96% | 96% |
| Damaraland mole rat (Fukomys damarensis) | 96% | 96% |
| Tibetan antelope (Pantholops hogsonii) | 96% | 96% |
| Blind mole rat (nannospalax galili) | 96% | 96% |
| Chinese hamster (Cricetulus griseus) | 95% | 95% |
| Guinea pig (Cavia procellus) | 95% | 95% |
| Golden hampster (Mesocricetus auratus) | 95% | 92% |
| Lesser Egyptian jerboa (Jaculus jaculus ) | 95% | 93% |
| Cow (Bos tauru) | 95% | 95% |

**Supplemental Table 1. Nucleotide sequence of the sTDP43-1 and -2 splice junctions are conserved in humans, non-human primates, and lesser mammals.**

| Species | Amino Acid Sequence | | | | | | | | | | | | | | | | | |
|---|---|---|---|---|---|---|---|---|---|---|---|---|---|---|---|---|---|---|
| *Reference Sequence* | V | H | L | I | S | N | V | Y | G | R | S | T | S | L | K | V | V | L |
| Chimpanzee (*Pan troglodytes*) | V | H | L | I | S | N | V | Y | G | R | S | T | S | L | K | V | V | L |
| Orangutan (*Pongo abelii*) | V | H | L | I | S | N | V | Y | G | R | S | T | S | L | K | V | V | L |
| Baboon (*Papio anubis*) | V | H | L | I | S | N | V | Y | G | R | S | T | S | L | K | V | V | L |
| Drill (*Mandrillus leucophaeus*) | V | H | L | I | S | N | V | Y | G | R | S | T | S | L | K | V | V | L |
| Black and white colobus (*Colobus angolensis palliatus*) | V | H | L | I | S | N | V | Y | G | R | S | T | S | L | K | V | V | L |
| Goat (*Capra hircus*) | V | H | L | I | S | N | V | Y | G | R | S | T | S | L | K | V | V | L |
| Cat (*Felis catus*) | V | H | L | I | S | N | V | Y | G | R | S | T | S | L | K | V | V | L |
| Alpine marmot (*Marmota marmota marmota*) | V | H | L | I | S | N | V | Y | G | R | S | T | S | L | K | V | V | L |
| Long-tailed chincilla (*Chinchilla lanigera*) | V | H | L | I | S | N | V | Y | G | R | S | T | S | L | K | V | V | L |
| Ground squirrel (*Ictidomys tridecemlineatus*) | V | H | L | I | S | N | V | Y | G | R | S | T | S | L | K | V | V | L |
| Kangaroo rat (*Dipodomys ordii*) | V | H | L | I | S | N | V | Y | G | R | S | T | S | L | K | V | V | L |
| Guinea pig (*Cavia porcellus*) | V | H | L | I | S | N | V | Y | G | R | S | T | S | L | K | V | V | L |
| Brown rat (*Rattus norvegicus*) | V | H | L | I | S | N | V | Y | G | R | S | T | S | L | K | V | V | L |
| Darama mole rat (*Fukomys damarensis*) | V | H | L | I | S | N | V | Y | G | R | S | T | S | L | K | V | V | L |
| Blind Mole Rat (*Nannospalax galili*) | V | H | L | I | S | N | V | Y | G | R | S | T | S | L | K | V | V | L |
| Golden hampster (*Mesocricetus auratus*) | V | H | L | I | S | N | V | Y | G | R | S | T | S | L | K | V | V | L |
| Deer mouse (*Peromyscus maniculatus bairdii*) | V | H | L | I | S | N | V | Y | G | R | S | T | S | L | K | V | V | L |
| House mouse (*Mus musculus*) | V | H | L | I | S | N | V | Y | G | R | S | T | S | L | K | V | V | L |
| Naked mole rat (*Heterocephalus glaber*) | V | H | L | I | S | N | V | F | G | R | S | T | S | L | K | V | V | L |
| Horse (*Equus calabus*) | V | H | L | M | S | N | V | Y | G | R | S | T | S | L | K | V | V | L |
| Tibetan antelope (*Pantholops hodgsonii*) | V | H | L | I | S | N | V | H | G | R | S | T | S | L | K | V | V | L |
| Chinese tree shrew (*Tupaia chinensis*) | V | H | L | I | S | N | V | S | G | R | S | T | S | L | K | V | V | L |

**Supplemental Table 2. Amino acid sequence of the sTDP43 C-terminal tail is conserved in humans, non-human primates, and lesser mammals.**

| Construct | Source | Complementary Oligomers | Sequence (5' to 3') |
|---|---|---|---|
| pUCM-CLYBL-NGN1-2-RFP | Gift from M. Ward | | |
| pLTC13-L1 | Gift from M. Ward | | |
| pLTC13-R1 | Gift from M. Ward | | |
| pX335-U6-Chimeric_BB-CBh-hSpCas9n(D10A) | Addgene (42335, donated by Feng Zhang) | | |
| pX330S-4 | Addgene (58780, donated by Feng Zhang) | | |
| pUCM-N-term-TARDBP-D2-HDR | Synthesized by Blue Heron, LLC | | |
| pX335-sgRNA-D2-TDP43-Upstream | This paper | Sense | GACCGTTCATATCTCTTTTCTCTTT |
| | | Antisense | AAACAAAGAGAAAAGAGATATGAAC |
| pX335-sgRNA-D2-TDP43-Downstream | This paper | Sense | CACCGGGGCTCATCGTTCTCATCTT |
| | | Antisense | AAACAAGATGAGAACGATGAGCCCC |
| pUCM-C-term-TARDBP-D2-HDR | Synthesized by Blue Heron, LLC | | |
| pX335-sgRNA-TDP43-D2-Upstream | This paper | Sense | CACCGGTTGGTTGGTATAGAATGG |
| | | Antisense | AAACCCATTCTATACCAACCAACC |
| pX335-sgRNA-TDP43-D2-Downstream | This paper | Sense | CACCGACCACTGCCCGACCCTGCAT |
| | | Antisense | AAACATGCAGGGTCGGGCAGTGGTC |

**Supplemental Table 3. Constructs and primer sequences used to generate iPSC lines**

| Antibody | Source | Catalog Number | Species | Dilution |
|---|---|---|---|---|
| Primary | | | | |
| Vglut1 | Synaptic Systems | 135303 | rabbit | 1:200 for ICC in iNeurons |
| Tuj1 | BioLegend | 801202 | mouse | 1:500 for ICC in iNeurons |
| N-term TDP43 | Sephton et al. 2011; Barmada et al. 2014 | NA | rabbit | 1:5000 for ICC and western |
| C-term TDP43 | Sephton et al. 2011; Barmada et al. 2014 | NA | rabbit | 1:5000 for ICC and western |
| sTDP43 | Custom-made from Genscript | NA | rabbit | 1:1000 for ICC and western |
| | | | | 1:500 for murine IHC |
| | | | | 1:250 for human IHC |
| Map2 | Milipore | MAB3418 | mouse | 1:1000 for ICC in iNeurons |
| GAPDH | Milipore | MAB374 | mouse | 1:1000 for ICC in iNeurons |
| GFAP | Abcam | AB53554 | goat | 1:500 for murine IHC |
| NeuN | Abcam | AB104225 | mouse | 1:500 for murine IHC |
| GFAP | Milipore | AB5541 | chicken | 1:500 for human IHC |
| NeuN | Milipore | MAB377 | mouse | 1:250 for human IHC |
| TDP43 | R&D Biosystems | MAB7778 | mouse | 1:1000 for western and ICC |
| | | | | 1:500 for ICC in iPSC-derived astrocytes |
| | | | | 1:250 for human IHC |
| GFAP | Sigma | C9205 | mouse | 1:500 for ICC in iPSC-derived astrocytes |
| Secondary | | | | |
| Alexa Fluor 488 | Life Technologies | A11034 | goat anti-rabbit | 1:500 |
| Alexa Fluor 594 | Life Technologies | A11032 | goat anti-mouse | 1:500 |
| Alexa Fluor 647 | Life Technologies | A31573 | donkey anti-rabbit | 1:500 |
| Alexa Fluor 647 | Life Technologies | A21449 | goat anti-chicken | 1:500 |
| LI-COR 680 | LI-COR Biosciences | 926-68072 | donkey anti-mouse | 1:5000 |
| LI-COR 800 | LI-COR Biosciences | 925-32213 | donkey anti-rabbit | 1:5000 |
| Alexa Fluor 488 | Life Technologies | AB150113 | goat anti-mouse | 1:500 |
| Alexa Fluor 647 | Life Technologies | AB150131 | donkey anti-goat | 1:500 |
| Alexa Fluor 568 | Life Technologies | AB175470 | donkey anti-rabbit | 1:500 |
| Alexa Fluor 647 | Thermo Fisher | A-21245 | goat anti-rabbit | 1:500 |
| Alexa Fluor IgG1 | Thermo Fisher | A-21121 | goat anti-mouse | 1:1000 |

### Supplemental Table 4. Antibodies

| Target | Location | Primer | Sequence (5' to 3') |
| --- | --- | --- | --- |
| ARC | *ARC* exon 2 | Forward | CCTGTACCAGACGCTCTACG |
| | | Reverse | GCAGGAAACGCTTGAGCTTG |
| Total TARDBP | *TARDBP* exon 1 and 2 | Forward | CTGCTTCGGTGTCCCTGTC |
| | | Reverse | TGGGCTCATCGTTCTCATCT |
| full-length (fl) TARDBP | *TARDBP* exon 6 stop codon | Forward | GTGGCTCTAATTCTGGTGCAG |
| | | Reverse | CACAACCCCACTGTCTACATT |
| sTDP43-1 | *sTDP43-1* splice donor | Forward | AGAAGTGGAAGATTTGGTGTTCA |
| | | Reverse | GCATGTAGACAGTATTCCTATGGC |
| sTDP43-2 | *sTDP43-2* splice donor | Forward | AGATTTGGTGGTAATCCAGTTCA |
| | | Reverse | GGCCTGTGATGCGTGATGA |
| CFTR minigene | *CFTR* exon 9 splice junction | Forward | CAACTTCAAGCTCCTAAGCCACTGC |
| | | Reverse | TAGGATCCGGTCACCAGGAAGTTGGTTAAATCA |

**Supplemental Table 5. Primers used in RT PCR and CFTR Assays**

| Plasmid | Source | Amplicon or Insert | Primer | Sequence (5' to 3') |
|---|---|---|---|---|
| pGW1-mApple | Barmada et al. 2014 | | | |
| pGW1-EGFP(1) | Arrasate et al. 2004 | | | |
| pGW1-TDP43-EGFP | Barmada et al. 2014 | | | |
| pGW1-sTDP43-EGFP | This paper | sTDP43 | NA | |
| pGW1-sTDP43(mNES)-EGFP | This paper | sTDP43(mNES) | Forward | CGCGGGCCCATGTCTGAATATATTCG |
| | | | Reverse | GCGACCGGTCGCAGCACTCCACCCCCTCCGCCGCTTCTTCCATAAAC |
| pGW1-EGFP(2) | This paper | EGFP | Forward | GCGAAGCTTGCCACCATGGTGAGCAAG |
| | | | Reverse | CGCGGTACCCTTGTACAGCTCGTCCAT |
| pGW1-EGFP-tail | This paper | tail | Forward | CGTTCATCTCATTTCAAATGTTTATGGAAGAAGCACTTCATTGAAAGTAGTGCTGTAAG |
| | | | Reverse | CTAGCTTACAGCACTACTTTCAATGAAGTGCTTCTTCCATAAACATTTGAAATGAGATGAACGGTA |
| pGW1-EGFP-tail(mNES) | This paper | tail(mNES) | Forward | CGTTCATCTCATTTCAAATGTTTATGGAAGAAGCGGCGGAGGGGGTGGAGTGCTGTAAG |
| | | | Reverse | CTAGCTTACAGCACTCCACCCCCTCCGCCGCTTCTTCCATAAACATTTGAAATGAGATGAACGGTA[illegible] |
| pGW1-EGFP-flTDP43 | This paper | TDP43 | Forward | GCG GGT ACC ATG TCT GAA TAT ATT CGG |
| | | | Reverse | CGC GCT AGC TTA TCC CCA GCC AGA AG |
| pGW1-EGFP-sTDP43 | This paper | sTDP43 | Forward | GCGGGTACCATGTCTGAATATATTCGG |
| | | | Reverse | CGCGCTAGCTTACAGCACTACTTTCAATG |
| pGW1-Halo | This paper | Halo | Forward | AAAAAA TCTAGA GCCACCATGGCAGAAATCGG |
| | | | Reverse | AAAAAA CCTGCAGG CTA GGAAATCTCGAGCGTCGACA |
| pGW1-flTDP43-Halo | This paper | TDP43, Halo | Forward | AAAAAA TCTAGA ATGGCAGAAATCGGTACTGG |
| | | | Reverse | AAAAAA CCTGCAGG CTA GGAAATCTCGAGCGTCGACA |
| pGW1-sTDP43-Halo | This paper | sTDP43 | Forward | GCG GCTAGC GCCACC ATGTCTGAATATATTCG |
| | | | Reverse | CGCACCGGTGGCAGCACTACTTTC |
| pCaggs-mCherry-TARDBP3'UTR (TDP43 autoregulation reporter) | This paper | TARDBP Exon 6 and 3'UTR | Forward | ATATTGTACATTGCGCAGTCTCTTTGTGGA |
| | | | Reverse | ATATGGCCGAGGCGGCCATCGTGTTTTTCCAGTAAGACTCCAGAC |
| pcDNA3.1-NLS-mCherry-NES (shuttle-RFP) | Addgene (72660, donated by B. Di Ventura and R. Eils) | | | |
| pFN21A HaloTag PUM2 RBD R6SYE | Gift from A. Goldstrohm | | | |
| pTB CFTR A455E TG13T5 (CFTR minigene) | Gift from Y. Ayala | | | |
| pGW1-CMV | Gift from S. Finkbeiner | | | |
| pCaggs-mCherry | Gift from S. Finkbeiner | | | |
| pSMART Lenti-shTARDBP (human) CAG-TurboRFP-VSVG | Dharmacon (V3SH11240-224779127) | | | |
| pSMART Lenti-NT shRNA CAG-TurboRFP-VSVG | Dharmacon (VSC11719) | | | |

### Supplemental Table 6. Source and Construction of Plasmid Vectors

| Name | Source | Identifiers | Additional Information | Application |
|---|---|---|---|---|
| HEK293T | ATCC | CRL-3216; RRID:CVCL_0063 | NA | Figures 6A-E and 8A, Supplemental Figures 4, 5, 6, 7D,E, 8, 11A |
| Neuro-2A | ATCC | CCL-131; 62278033 | NA | Supplemental Figure 11B-E |
| Primary neurons | University of Michigan Unit for Laboratory Animal Medicine | NA | Derived as described in the Supplemental Materials and Methods | Figures 4, 5, 6F-H, Supplemental Figures 1, 3, 5A, 7B,C, 11F |
| Integrated gCaMP6f iPSC line | Michael Uhler, University of Michigan | NA | Stable integration of Ngn2 and gCaMP6f into a commercially available line at WiCell Research Resources (RRID: CVCL_9773) | Figure 1D-G |
| iPSC Line #1 | University of Michigan ALS Repository | NA | Stable integration of Ngn2; Insertion of Dendra2 5' or 3' of TARDBP | Figure 2B-E, Supplemental Figure 2 |
| iPSC Line #2 | University of Michigan ALS Repository | NA | Stable integration of Ngn2 | Figures 1C,H-K, 3C,D, and 8B,C |
| iPSC Line #3 | Thermo Fisher | Catalog Number A18945 | Human episomal iPSC line | Supplemental Figure 13 |

### Supplemental Table 7. Source and Description of All Cell Lines

15. D'Erchia AM, et al. Massive transcriptome sequencing of human spinal cord tissues

provides new insights into motor neuron degeneration in ALS. *Sci Rep*. 2017;7(1):10046.
